# Bacterial single-cell RNA sequencing captures biofilm transcriptional heterogeneity and differential responses to immune pressure

**DOI:** 10.1101/2024.06.28.601229

**Authors:** Lee E. Korshoj, Tammy Kielian

**Affiliations:** Department of Pathology, Microbiology, and Immunology, University of Nebraska Medical Center, Omaha, NE, USA

**Keywords:** bacterial single-cell RNA sequencing, biofilm, infection, bacterial heterogeneity, bacterial transcriptomics, immune response, leukocytes, *Staphylococcus aureus*, split-pool barcoding

## Abstract

Biofilm formation is an important mechanism of survival and persistence for many bacterial pathogens. These multicellular communities contain subpopulations of cells that display vast metabolic and transcriptional diversity along with high recalcitrance to antibiotics and host immune defenses. Investigating the complex heterogeneity within biofilm has been hindered by the lack of a sensitive and high-throughput method to assess stochastic transcriptional activity and regulation between bacterial subpopulations, which requires single-cell resolution. We have developed an optimized bacterial single-cell RNA sequencing method, BaSSSh-seq, to study *Staphylococcus aureus* diversity during biofilm growth and transcriptional adaptations following immune cell exposure. We validated the ability of BaSSSh-seq to capture extensive transcriptional heterogeneity during biofilm compared to planktonic growth. Application of new computational tools revealed transcriptional regulatory networks across the heterogeneous biofilm subpopulations and identification of gene sets that were associated with a trajectory from planktonic to biofilm growth. BaSSSh-seq also detected alterations in biofilm metabolism, stress response, and virulence that were tailored to distinct immune cell populations. This work provides an innovative platform to explore biofilm dynamics at single-cell resolution, unlocking the potential for identifying biofilm adaptations to environmental signals and immune pressure.

## INTRODUCTION

Bacterial infections represent a pervasive clinical problem that is increasingly complicated by the emergence of multidrug-resistant (MDR) strains, recognized as one of the greatest threats to human health worldwide.^1–3^ One successful bacterial pathogen typified by MDR is *Staphylococcus aureus* (*S. aureus*).^4^ While a commensal in nearly one-third of the human population, *S. aureus* is transmitted across both hospital and community settings as a leading cause of post-surgical infection, skin and soft tissue infection, bacteremia, endocarditis, osteomyelitis, and medical device-associated infection.^5^ In addition to the large arsenal of immune evasion molecules and antibiotic resistance genes encoded by *S. aureus*, a hallmark of this pathogen is its propensity for biofilm formation.^5,6^ Biofilm is a key mechanism for survival and persistence in the infected host, leading to significant morbidity and mortality not only for *S. aureus*, but also other MDR pathogens including *Escherichia coli*, *Klebsiella pneumonia*, and *Pseudomonas aeruginosa*.^7^ It has been estimated that approximately 65% of nosocomial infections are associated with biofilm formation.^8^ Encased in an extracellular matrix comprised of polysaccharides, proteins, and nucleic acids, the multicellular biofilm community is highly recalcitrant to antibiotics and the host immune system.^6–8^ A combination of bulk transcriptomics, bacterial mutants, and fluorescent reporter strains have been employed to identify metabolically and transcriptionally diverse subpopulations of bacterial cells within biofilm that have differing roles in surface attachment, dispersal and dissemination, stress-response, host defense, and persistence.^6,7^ Understanding these communities has been hindered by lack of a high-throughput method to simultaneously measure the complex and stochastic interactions between distinct bacterial subpopulations.

Single-cell RNA sequencing (scRNA-seq) is widely used for transcriptional profiling of eukaryotic cells within a heterogeneous sample.^9^ It has been applied to assess immune response dynamics during bacterial infection, including biofilm, identifying transcriptional changes in leukocyte metabolism, reactive oxygen species (ROS) production, and inflammatory mediator signaling specific to each immune cell type.^10–14^ However, the use of scRNA-seq has traditionally been limited in prokaryotes based on the short half-life and low abundance of mRNA, lack of polyadenylated transcripts, and complex cell wall characteristics.^15–17^ As a result, bulk RNA-seq methods have primarily been used to study bacterial pathogens and biofilm communities. However, bulk methods fail to capture heterogeneity and underrepresented populations altogether. A single-cell approach is necessary for a complete transcriptional landscape of biofilm heterogeneity and how biofilm is affected in response to distinct immune pressures, a critical step towards identifying novel anti-biofilm strategies.

Only recently have bacterial scRNA-seq methodologies been described, each employing unique protocol variations with respective pros and cons.^18–24^ One major area of variation between described methods is how individual cells are labeled with distinct oligonucleotide barcode sequences for identification, with methods broadly separating into plate- and microfluidics-based barcoding approaches. Plate-based approaches have utilized standard 96- or 384-well plates that impose inherent limitations on cell numbers,^18,20,23^ while microfluidics-based approaches permit acquisition of increased cell numbers but require adaptation of costly commercial instrumentation.^21,22,24^ Another technique employed fluorescence-activated cell sorting (FACS) for bacterial cell separation and identification, but cell yields were limited to a few hundred.^19^ A second major area of methodological variation is RNA capture, with most methods utilizing random hybridization or mRNA-targeted probes. The use of targeted mRNA probes requires prior knowledge of the genome and desired targets, effectively limiting the number of genes analyzed,^22^ while random hybridization provides unbiased insights into all possible genes but results in an overabundance of rRNA reads (i.e., >90%).^18,20,21,23,24^ Initial studies with random RNA hybridization omitted rRNA depletion, whereas more recent reports successfully incorporated rRNA depletion with Cas9 or RNase H.^21,24^ In all published bacterial scRNA-seq methods to date, studies were limited to planktonic organisms and focused on proof-of-concept feasibility of the approach. Several reports examined transcriptional changes between different planktonic growth states,^18,20^ whereas others observed transcriptional variation arise in planktonic culture upon treatment with antibiotics or other stimuli.^21,22,24^

Here, we present an advanced method and application of bacterial scRNA-seq to explore the heterogeneity of complex biofilm communities and transcriptional adaptations in response to immune cell challenge. Our technique, termed BaSSSh-seq (<u>ba</u>cterial single-cell RNA sequencing with <u>s</u>plit-pool barcoding, <u>s</u>econd strand synthesis, and <u>s</u>ubtractive <u>h</u>ybridization), employs an optimized protocol for RNA capture from bacterial cells with low metabolic activity, as seen in biofilm. BaSSSh-seq uses plate-based split-pool barcoding to label individual cells, without the need for sophisticated commercial equipment.^25–28^ Random hexamers are used for unbiased RNA capture during barcoding. Additionally, second strand synthesis replaces the highly inefficient process of template switching to generate cDNA libraries,^29^ and an enzyme-free rRNA depletion method based on subtractive hybridization is used to significantly reduce rRNA contamination.^30^ Through reduced enzyme usage and rRNA contamination, costs are decreased while concurrently increasing sequencing depth. This concept is important for bacterial scRNA-seq given the inherent sparseness of cellular mRNA. We established that diversity can be captured from bacterial cells with low metabolic and transcriptional activity within biofilm and coupled this with innovative computational assessments for identifying transcriptional heterogeneity and dynamics.

We applied BaSSSh-seq to study unique transcriptional signatures that differentiate *S. aureus* biofilm from planktonic growth and how biofilm alters its transcriptional profile in response to immune pressure, elevating bacterial scRNA-seq from proof-of-concept demonstrations to address complex biological interactions. An initial comparison of biofilm vs. planktonic growth demonstrated the ability to capture transcriptional heterogeneity within biofilm and validated the BaSSSh-seq methodology through extensive consistency with literature and experimental observations. We then explored biofilm transcriptional alterations in response to immune pressure by applying BaSSSh-seq to biofilm after direct co-culture with three major leukocyte populations that have well-documented roles in *S. aureus* infection: macrophages (MΦs), neutrophils (PMNs), and granulocytic myeloid-derived suppressor cells (G-MDSCs).^31–33^ Within the transcriptionally diverse subpopulations of biofilm, differential responses to each leukocyte population were observed. We further developed a powerful analytical pipeline using a combination of unique computational assessments and existing bioinformatics packages for an enhanced multi-level visualization of biofilm transcription. Through integration of iModulon analyses, we achieved a high-level assessment of transcriptional regulatory networks across biofilm subpopulations in addition to gene-level characterization.^34–36^ Likewise, trajectory analysis was used to identify transcriptional dynamics between *S. aureus* growth states and activation upon immune pressure.^37^ Together, BaSSSh-seq provides the opportunity for studying biofilm growth dynamics and interactions with the immune system at a new level of resolution, promoting enhanced understanding of biofilm pathogenesis and the potential for rational design of new therapeutic strategies.

## RESULTS

### BaSSSh-seq enables bacterial scRNA-seq of biofilm and incorporates rRNA depletion

We employed split-pool barcoding to capture and label RNA transcripts (Figure 1A), a technique originally described in eukaryotic cells and recently applied to prokaryotes.^18,20,25–28^ Split-pool barcoding attaches a cell-specific combination of three oligonucleotide barcodes to RNA transcripts. Barcoding is performed in fixed and permeabilized bacteria over three rounds consisting of an initial reverse transcription reaction where the RNA is captured, followed by two ligation reactions, interspersed with pooling and mixing steps. In our optimized implementation of split-pool barcoding, random hexamers were used for RNA capture during reverse transcription along with blocking unreacted barcodes with a set of complementary oligos during pooling to prevent non-specific and erroneous barcode ligations.^18^ We also filtered, vortexed, and briefly sonicated cells between each barcoding step, which was previously shown to decrease the doublet rate.^20^ Advantages of split-pool barcoding include its feasibility and cost, requiring only standard laboratory equipment. Other renditions of bacterial scRNA-seq have adapted commercial microfluidic instruments for cellular barcoding, requiring access to costly specialized equipment and reagents.^21,22,24^

**Figure 1.**
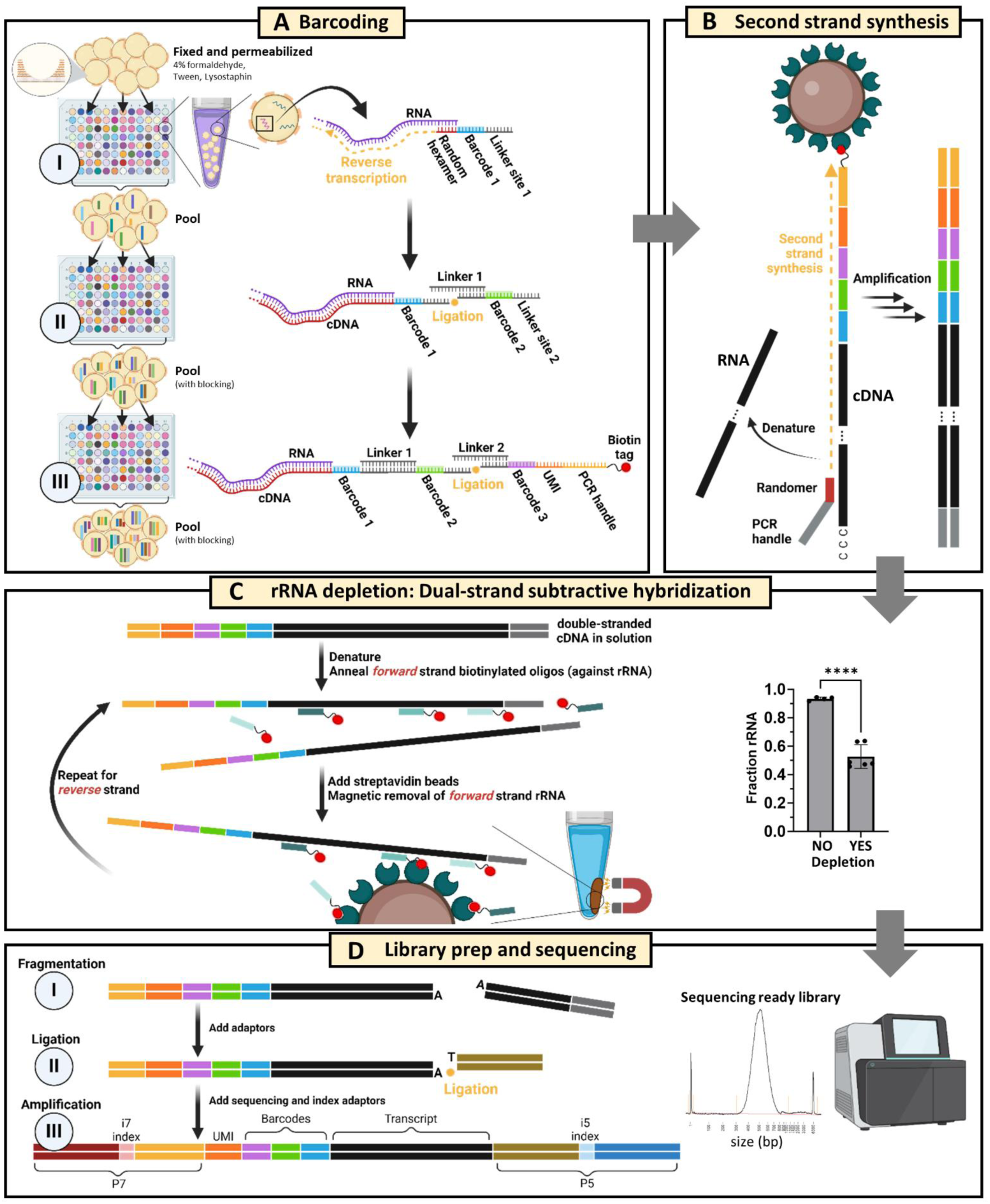
BaSSSh-seq enables bacterial scRNA-seq of biofilm and incorporates rRNA depletion. (A) Split-pool barcoding attaches a combination of three barcodes to intracellular RNA transcripts of fixed and permeabilized cells. The 5’ end of the terminal barcode oligo also includes a UMI, PCR handle, and biotin tag. (B) Following lysis, streptavidin magnetic beads are used to purify captured transcripts. Then double-stranded cDNA is synthesized via random primer second strand synthesis and PCR amplification. (C) Substantial rRNA depletion is performed using an enzyme-free dual-strand subtractive hybridization technique, where biotin-tagged oligos specific to 5S, 16S, and 23S rRNA fragments are annealed to each cDNA strand and magnetically removed with streptavidin beads. The rRNA content can be lowered from >90% to <50%. ****, *p*-value <0.0001 by unpaired t test (4 biological replicates with no depletion, 6 biological replicates with depletion). (D) Libraries are constructed for Illumina sequencing through fragmentation, ligation, and amplification to generate constructs containing P5/P7 ends with unique i5/i7 index combinations. Schematics created in BioRender.

Following barcoding, cells were lysed, and captured transcripts were purified with streptavidin magnetic beads leveraging a biotin tag on the 5’-end of the terminal barcode oligo. For generation of double-stranded cDNA, a second PCR handle is required. A typical technique used for this is template switching, which exploits the terminal transferase activity of certain reverse transcriptase enzymes to anneal and synthesize the required PCR handle.^38^ Reliance on a short ∼3-nucleotide sequence for annealing is highly inefficient and leads to significant transcript loss. Additional complications include concatamerization of the switching oligo when template concentrations are very low^39^, which was observed for bacterial RNA samples (Figure S1A). Therefore, we incorporated a random primed second strand synthesis step (Figure 1B) as recently described for eukaryotic scRNA-seq, which significantly improved transcript capture compared to template switching.^29^

While random hexamer capture of RNA provides an unbiased survey of cellular transcripts, it leads to an overabundance of rRNA that can account for >90% of total sequencing reads, which we observed during protocol optimization and is consistent with known rRNA abundance in bacteria.^18,20,21,30^ Initial permutations of bacterial scRNA-seq omitted rRNA depletion, largely due to the difficulties in translating applicable depletion techniques from bulk RNA-seq to the in-cell reactions necessary for single-cell barcoding. Subsequent permutations employed RNase H and Cas9 methods for rRNA depletion prior to barcoding, reducing levels by approximately 50%.^21,24^ While a significant reduction, these procedures rely on additional enzymatic steps performed on fixed and permeabilized cells, potentially leading to substantial cell loss,^21^ which we also observed in initial studies on cells prior to barcoding. When working with precious samples at low cell numbers, any losses can negatively impact or bias results. Therefore, we applied an enzyme-free rRNA depletion process to our double-stranded cDNA pool to reduce cell loss (Figure 1C). This strategy uses subtractive hybridization, where short biotinylated oligos are annealed to rRNA-derived cDNA sequences and removed using magnetic beads.^30^ Applied to double-stranded cDNA, the process is conducted on both forward and reverse strands. With this approach, rRNA levels were reduced to below 50%, consistent with other bacterial scRNA-seq methods but with the advantage of less cell loss from additional enzymatic steps.

After rRNA depletion, cDNA was fragmented to an optimal sequencing size (400-700 bp), ligated with short adaptors, and amplified to yield Illumina-compatible sequencing libraries containing P5 and P7 regions with dual indices (Figure 1D). Library constructs contained a UMI for consolidating PCR duplicates and barcodes in read 2, and the transcript in read 1. BaSSSh-seq was shown to faithfully capture bacterial transcriptomic profiles by comparing results with a traditional bulk RNA-seq dataset of *S. aureus* biofilm previously generated in our laboratory that yielded statistically significant Pearson correlations (Figure S2).^40^ Given the intrinsic heterogeneity of biofilm, and variability in growth and sampling over the timescales between the two datasets, this provides strong validation of BaSSSh-seq fidelity. For our *S. aureus* samples, BaSSSh-seq captured an average of 12–60 mRNA reads per cell, consistent with other bacterial scRNA-seq methods applied to *S. aureus* under planktonic growth conditions.^18^ Considering that our analysis focused on biofilm, which is known to contain less metabolically and transcriptionally active cells,^6^ this highlights the importance of our optimized protocol improvements. Although the mRNA counts achieved for *S. aureus* biofilm are less than those reported for another widely studied gram-positive pathogen, *Bacillus subtilis*, where 200–300 mRNA reads per cell were captured during planktonic growth,^20,22^ this discrepancy can be explained by its larger cell volume (∼4-6X) compared to *S. aureus*.^41–43^ Studies from eukaryotic scRNA-seq have established that low-coverage sequencing is sufficient to fully capture sample heterogeneity within a large number of cells.^44–47^ Quality control measures throughout the BaSSSh-seq process are presented in the Methods and Figure S1B-E.

### Biofilm growth is marked by extensive transcriptional heterogeneity and decreased metabolic gene expression at the single-cell level

Our first examination into *S. aureus* biofilm transcriptional complexity using BaSSSh-seq was with direct comparison to planktonic growth (Figure 2A). As previously mentioned, earlier studies using bacterial scRNA-seq focused on planktonic growth and reported that unperturbed planktonic cultures are largely homogeneous.^18,20–22^ Biofilm has been characterized to contain a heterogeneous population of cells with varying microstructural attributes that experience coordinated physiological changes throughout development.^48–52^ Full mechanistic insights into signaling within the biofilm network remains elusive as our current understanding has principally relied on time-lapse microscopy with limited transcriptional reporter panels.^53–55^ Comparisons between biofilm and planktonic growth have been explored in multiple studies over the past two decades with bulk transcriptomic or proteomic techniques.^56–59^ Our BaSSSh-seq platform provides unprecedented resolution for this growth condition comparison, and the potential for deeper mechanistic understanding of transcriptional signaling within diverse biofilm communities. Cells from biofilm and planktonic samples were collected simultaneously and fixed overnight before permeabilization the following day under identical conditions. For the first round of barcoding, cells from biofilm and planktonic cultures were processed separately so sample origin could be identified post-sequencing. Samples were combined for the second and third rounds of barcoding and processed together for sequencing. The planktonic culture yielded more cells with higher amounts of captured mRNA compared to biofilm (Figure S3A-C). This was expected based on the known differences in cellular activity during exponential growth of planktonic bacteria compared to biofilm where many organisms display a metabolically dormant phenotype.^20,59^ To ensure equal assessment with comparable cell numbers, sequenced cells for biofilm and planktonic samples were filtered at 7 and 28 non-rRNA reads per cell, respectively, which resulted in similar cell numbers (biofilm n=3,680 and planktonic n=4,231, Figure S3A) for downstream bioinformatic analysis.

**Figure 2.**
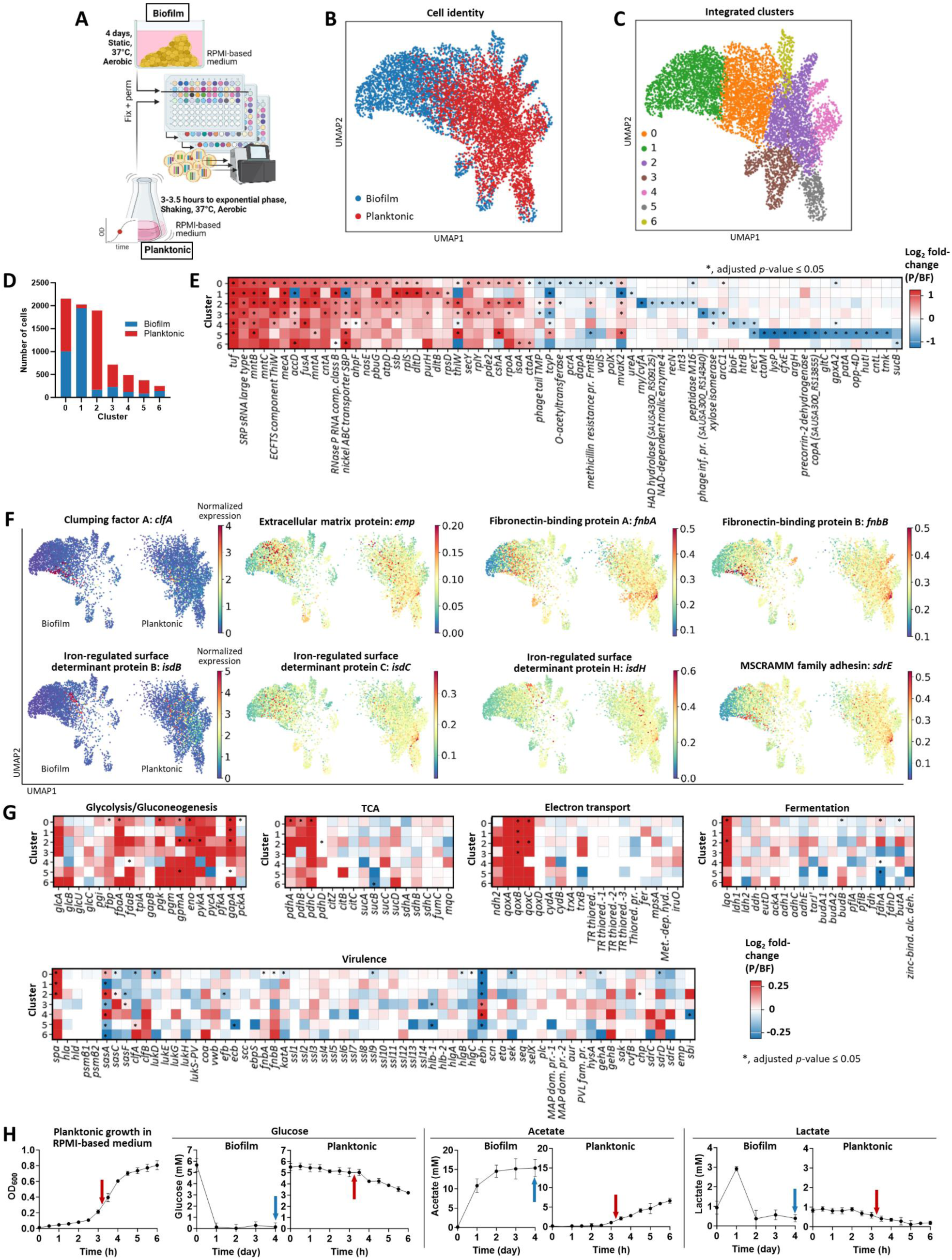
Biofilm growth is marked by extensive transcriptional heterogeneity and decreased metabolic gene expression at the single-cell level. (A) *S. aureus* biofilm was grown for 4 days under static conditions with daily medium replenishment, and planktonic culture was grown to exponential phase between 3-3.5 h with shaking at 250 rpm. Both biofilm and planktonic samples were grown in identical RPMI-based medium with aerobic incubation at 37°C. Cells from biofilm and planktonic samples were fixed overnight before permeabilization under identical conditions. Cells from biofilm and planktonic cultures were separate for the first round of barcoding, then combined for the second and third rounds. The combined samples were processed through to sequencing. (B and C) UMAP plots of the integrated biofilm and planktonic samples depicting (B) sample origin and (C) subpopulations identified with the Leiden algorithm. (D) Distribution of biofilm and planktonic cells across each cluster. (E) Marker genes specific to planktonic and biofilm growth across all clusters, represented as log_2_ fold-change of planktonic (P) / biofilm (BF). Red signifies upregulation in planktonic and blue signifies upregulation in biofilm. SRP = signal recognition particle, ECFTS = energy coupling factor transporter S, SBP = substrate-binding protein, TPM = tape measure protein. (F) Expression of exoproteome-associated genes overlaid on UMAP plots, separated by biofilm and planktonic. Color represents normalized expression level on a per cell basis. (G) Comparisons of metabolic and virulence factor gene expression between planktonic and biofilm growth, represented as log_2_ fold-change of planktonic (P) / biofilm (BF). Red signifies upregulation in planktonic and blue signifies upregulation in biofilm. (H) Quantification of glucose, acetate, and lactate in culture supernatants collected during biofilm and planktonic growth. The planktonic growth curve in RPMI-based medium is also shown on the left. Arrows indicate the time of sample collection.

Clustering analysis using uniform manifold approximation and projection (UMAP) was performed on the integrated biofilm and planktonic datasets after batch balanced k nearest neighbors (BBKNN) alignment.^60^ UMAP visualization indeed revealed greater spatial distribution of biofilm cells, reflecting enhanced transcriptional diversity (Figure 2B). In total, 7 transcriptionally unique subpopulations, or clusters, were identified across the integrated biofilm and planktonic dataset (Figure 2C and Figure S3D). The largest cluster (cluster 0), was most equally distributed between biofilm and planktonic cells (Figure 2D). The remaining clusters were largely biased towards biofilm or planktonic cells, reflecting the intrinsic transcriptional differences between the growth states. Differential gene expression was performed across all clusters using the MAST algorithm.^61^ This hurdle model is important to account for the large number of dropouts, or non-detected genes in scRNA-seq datasets, which was observed with the biofilm sample given the low metabolic activity of some bacterial subpopulations. A known caveat of MAST is that log_2_ fold-change values can be small; therefore, minor differences cannot be disregarded as insignificant. Each cluster contained a set of marker genes for biofilm and planktonic growth, a subset of which are detailed in Figure 2E. Overall, upregulated genes within the planktonic growth state were uniformly increased across the majority of clusters, whereas most upregulated genes within biofilm were unique to specific clusters, strongly highlighting its transcriptional heterogeneity.

During planktonic growth, several genes related to transcriptional and translational activity were more highly expressed compared to biofilm across the majority of clusters, including RNA polymerase subunit alpha (*rpoA*), elongation factors (*tuf*, *fusA*), and ribosomal proteins (*rplS*, *rplY, rpsD*).^62–64^ Upregulation of genes encoding an ATP synthase (*atpD*) and primary heme A component of terminal oxidases (*ctaA*) suggest increased respiration in planktonic cells.^65,66^ Heightened expression of genes for a single-stranded DNA-binding protein necessary for DNA replication (*ssb*), a regulator of secondary messenger cyclic di-adenosine monophosphate (c-di-AMP) (*pde2*), a DEAD-box RNA helicase with known control over *agr*-mediated quorum sensing (*cshA*), and a housekeeping protein (*isaA*) are indicative of cellular division and environmental sensing associated with *S. aureus* exponential growth.^67–71^ Planktonic cells also upregulated manganese transporters (*mntABC*) reported to combat oxidative stress generated from heightened respiratory activity during aerobic growth, and a set of lipoteichoic acid-associated genes for D-alanylation (*dltD, dltB*) that are linked to increased cellular fitness.^72–74^

As mentioned, upregulated genes within biofilm were primarily limited to specific clusters, reflecting increased heterogeneity. An interesting observation was that biofilm clusters were enriched for genes associated with genetic variation, including plasmid replication (*pcrA*), DNA repair (*recN*), integrase activity (*int3*), and recombinase activity (*recT*).^75–80^ Biofilm also expressed several phage protein genes (cataloged as ‘*phage tail tape measure protein*’ and ‘*phage infection protein’*). While likely remnants of previous phage insertion, phage activity has been postulated to promote bacterial persistence and survival during biofilm maturation and remodeling.^75^ Upregulation of genes for arginine and lysine biosynthesis (*argH* and *dapA*), cysteine transport (*tcyP*), histidine metabolism (*hutI*), oligopeptide transport (*opp-4D*), and glutamate regulation (*gltC*) suggest reliance on amino acids for a range of cellular processes within biofilm since these genes are linked to nutrition, signaling, and virulence.^81–87^ Further, nutrient limitation and stress were evident in biofilm by increased expression of genes for biotin synthesis (*bioF*), copper transport (*copA*), and urease (*ureA*) that is important for pH regulation within biofilm.^59,88,89^ Increased RNase Y (*rny/cvfA*) levels were also seen in biofilm, which has been shown to tightly control mRNA expression for coordinated virulence gene activation.^90^

*S. aureus* exoproteome-associated genes have well-characterized roles in virulence and biofilm formation.^91–96^ To examine their expression patterns across biofilm and planktonic growth states, gene expression was overlaid onto the UMAP space with the MAGIC imputation algorithm to remove noise obscuring underlying expression patterns, due to the inherent dropouts in scRNA-seq datasets.^97^ Many exoproteome-associated genes displayed heighted expression in biofilm cells (Figure 2F). Interestingly, some appeared more diffuse throughout biofilm (*emp*, *fnbA*, *isdC*), while others displayed more concentrated expression patterns within specific clusters (*clfA*, *isdB*, *sdrE*). The expression of these genes during planktonic growth was concentrated to a single cluster that exhibited widespread gene induction, which we show later to be the most transcriptionally active cells within the planktonic culture.

Next, metabolic and virulence factor gene expression was compared between *S. aureus* growth states. For metabolic assessments, differential expression was performed between planktonic and biofilm cells across all clusters for genes in major metabolic pathways including glycolysis/gluconeogenesis, tricarboxylic acid (TCA) cycle, electron transport, and fermentation (Figure 2G). Glycolysis showed preferential upregulation in planktonic cells across all clusters compared to biofilm. Additionally, planktonic bacteria displayed increased expression of genes encoding the pyruvate dehydrogenase multienzyme complex (*pdhABCD*) that converts pyruvate to acetyl-CoA and acetate kinase (*ackA*) for acetate and ATP generation. This is consistent with known mechanisms of catabolite control protein A (CcpA) regulation under aerobic conditions and glucose availability, where the expression of glycolytic genes is increased while acetyl-CoA is converted to acetate and the TCA cycle is suppressed.^98^ L-lactate-quinone oxidoreductase (*lqo*), which converts L-lactate to pyruvate for downstream ATP generation, was also upregulated in planktonic cells, suggestive of moderate lactate utilization for respiration and growth.^99–101^ Planktonic cells further showed large induction of terminal oxidase genes (*qoxABCD*) required for respiration, consistent with the identified marker genes in Figure 2E.^102^ Expression of fermentative genes trended higher in biofilm, especially those related to formate metabolism (*pflAB* and *fdh*) that is important for biofilm structure and persistence.^40,103^ These changes in metabolic gene expression aligned with extracellular glucose, acetate, and lactate concentrations in supernatants from biofilm and planktonic cultures as a validation of the BaSSSh-seq system (Figure 2H). Compared to biofilm, exponential phase planktonic culture contained higher glucose levels that were progressively depleted, supporting the heightened expression of glycolytic genes. Acetate production in planktonic culture actively increased while the levels in mature biofilm plateaued, supporting the increased expression of *pdhABCD* and *ackA* in planktonic cells. Finally, lactate originating from the RPMI-based medium (Figure S3E) was actively consumed during planktonic growth whereas levels remained stable in the mature biofilm during later stage growth, supporting the observed upregulation of *lqo*. Virulence genes trended towards higher expression in biofilm compared to planktonic bacteria (Figure 2G), with the exception of Protein A (*spa*).^104^ One of the most highly upregulated genes within biofilm was for a giant surface protein (*ebh*) with noted roles in regulating *S. aureus* clumping, virulence, osmolarity, surface attachment, and biofilm formation.^43,105^ Another gene involved in surface attachment (*sasA*) was also increased in distinct biofilm clusters.^106^ Collectively, this first comparison of biofilm and planktonic growth at the single-cell level illustrated the powerful capabilities of scRNA-seq to capture transcriptional heterogeneity within biofilm while at the same time validating the robustness of the BaSSSh-seq methodology through corroboration of observed transcriptional patterns with the published literature.

### Transcriptional regulation follows a trajectory from planktonic to biofilm growth

While the prior analyses uncovered alterations between biofilm and planktonic growth states at the gene expression level (Figure 2), this did not provide insights into large-scale transcriptional regulation. Several computational tools exist for conducting pathway analysis for eukaryotic scRNA-seq, where *a priori* defined gene sets are assessed based on shared biological function.^107^ Similar tools for prokaryotes are designed for higher density data produced by bulk RNA-seq and cannot be functionally translated to sparser single-cell datasets or contain an extraneous number of pathways confounding interpretation.^108,109^ Recently, the concept of independent component analysis (ICA) has been applied to identify co-regulated, independently modulated gene sets (iModulons) within bacterial transcriptomes in an effort to understand the complex crosstalk between metabolism and gene regulation.^34–36^ For *S. aureus*, ICA was applied to >300 bulk RNA-seq datasets across a range of conditions yielding 76 iModulons.^34^ These iModulons were further condensed into 10 groups representing the transcriptional regulatory network. We integrated the ICA-determined gene sets from the *S. aureus* iModulonDB database into our BaSSSh-seq study as a new prokaryotic pathway analysis tool (Figure 3A).^36^

**Figure 3.**
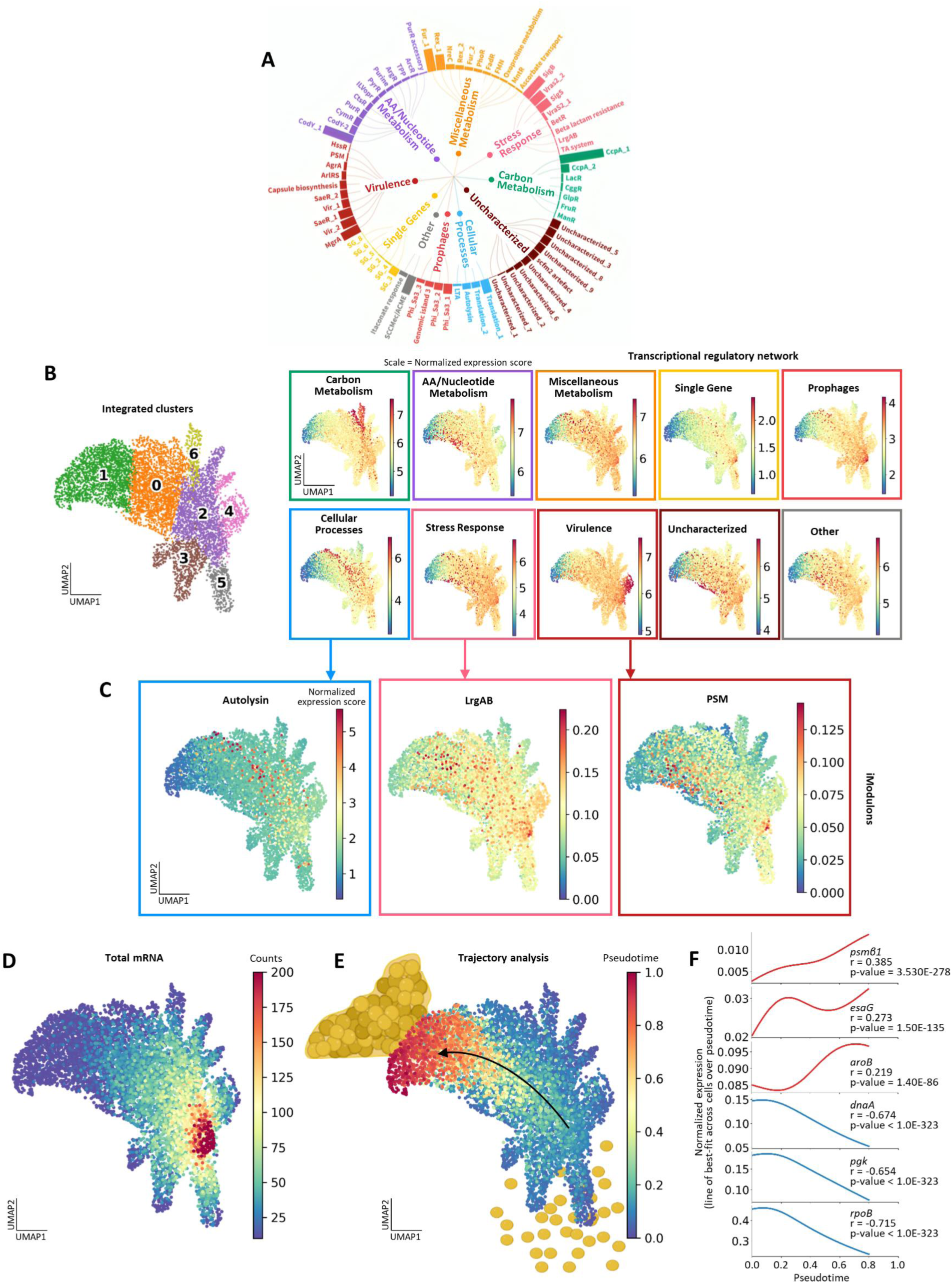
Transcriptional regulation follows a trajectory from planktonic to biofilm growth. (A) The *S. aureus* transcriptional regulatory network. Regulatory categories are comprised of sets of iModulons, where each iModulon is comprised of an independently modulated set of genes. The size of the bars corresponds to the number of genes in the iModulon. (B) Transcriptional regulatory categories are overlaid on UMAP plots (right). Color represents normalized expression score per cell. For reference, the cluster identities (left) are shown as defined in Figure 2C. (C) Autolysin, LrgAB, and PSM iModulon expression scores, linked to their respective transcriptional regulatory category. (D) Overlay of total mRNA counts on the UMAP plot, where color represents mRNA counts per cell. (E) Trajectory analysis (Palantir algorithm) identified a differentiation pathway through the integrated biofilm and planktonic samples, moving through pseudotime from planktonic to biofilm. (F) Subsets of genes display positive and negative correlation with the pseudotime trajectory. Additional genes are listed in Tables S1-S2.

Visualization of the *S. aureus* transcriptional regulatory network by overlaying expression scores onto the UMAP provided a meaningful coarse-grained view of metabolism, virulence, stress, and other cellular processes across biofilm and planktonic growth states (Figure 3B, right). Activity within specific regulatory categories can be linked to previously defined clusters reflecting planktonic or biofilm growth (Figure 3B, left, as defined in Figure 2C). Overall, planktonic cells exhibited the highest expression scores for most regulatory categories, reflecting increased activity (Figure 3B). In contrast, a region of cells in cluster 1 were inactive or dormant across all the regulatory categories, which mainly represent biofilm. Individual iModulon expression scores were overlaid on the UMAP for a more granular view (Figure 3C). The Autolysin, LrgAB, and PSM iModulons were selected based on the known associations of their respective genes with biofilm formation.^110,111^ Interestingly, these iModulons were preferentially increased in biofilm cells and progressed seemingly along a path through the UMAP space from planktonic towards biofilm growth (Figure 3C).

To further explore this relationship, global transcriptional activity was visualized by overlaying total mRNA counts onto the UMAP (Figure 3D). Planktonic cells, particularly within cluster 2 were most transcriptionally active, while biofilm cells in cluster 1 were least active. Trajectory analysis was then performed using the Palantir algorithm.^37^ Adapted from eukaryotic scRNA-seq, trajectory algorithms identify paths of differentiation through a dataset by quantifying divergences in gene expression between nearest-neighbor cells. Applied to our integrated biofilm and planktonic dataset, Palantir identified a trajectory from planktonic cells to biofilm, terminating at the most inactive or dormant cluster of biofilm cells (Figure 3E and Figure S4A). The trajectory over pseudotime largely followed the patterns of iModulon expression (Figure 3C). Further, a Pearson correlation analysis was conducted for each gene over pseudotime to identify genes positively and negatively correlated with the trajectory (Figure 3F). A subset of genes positively correlated with biofilm trajectory included phenol-soluble modulin (*psmβ1*), nuclease toxin system (*esaG*), and menaquinone biosynthesis (*aroB*).^111–113^ Genes negatively correlated with pseudotime were associated with replication initiation (*dnaA*), glycolysis (*pgk*), and transcription (*rpoB*).^114–116^ Comprehensive lists of positively and negatively correlated genes are provided in Tables S1-S2. The ability to quantify, visualize, and correlate transcriptional regulation across heterogeneous subpopulations of bacteria and growth trajectories provides a new level of resolution towards advancing our understanding of the mechanisms contributing to biofilm formation and persistence.

### Biofilm shows coordinated transcriptional regulation

After comparing biofilm and planktonic growth and demonstrating the ability to visualize regulatory dynamics across heterogeneous bacterial populations and growth states, we next focused on deeper analysis of biofilm heterogeneity and how this is altered in response to immune cell exposure. Our laboratory and others have studied the immune response to *S. aureus* biofilm infection in several animal models and humans.^31–33,117–121^ The major leukocyte infiltrates associated with biofilm include MΦs, G-MDSCs, and PMNs, which are insufficient at clearing infection, which remains chronic.^31,32^ These immune cell populations exhibit different metabolic, phagocytic, epigenetic, and transcriptional responses during *S. aureus* biofilm infection.^117–119^ However, little is known about how biofilm adapts to each leukocyte type, especially at the single-cell level.

To investigate this question, BaSSSh-seq was applied to *S. aureus* biofilm directly co-cultured with MΦs, G-MDSCs, or PMNs (Figure 4A). After co-culture, immune cells were lysed to prevent downstream RNA contamination. The bacterial cells from biofilm control (no immune cells) and the co-cultured biofilms were fixed overnight before permeabilization. Bacteria from each respective biofilm sample were barcoded separately for the first round, combined for the second and third rounds, and processed for sequencing. Sequencing reads were only aligned to the *S. aureus* transcriptome as a second means of preventing any potential eukaryotic RNA contamination. Sequenced bacteria were filtered at 15 non-rRNA reads per cell, which resulted in similar cell numbers per treatment (4,500 – 6,000 cells, Figure S5). UMAP clustering of the biofilm control yielded 7 transcriptional subpopulations (Figure 4B). Independent UMAP clustering of co-cultured biofilms led to unique patterns (Figure 4C and Figure S6), suggesting that *S. aureus* tailors its transcriptional response to each immune cell type, which will be described in more detail below.

**Figure 4.**
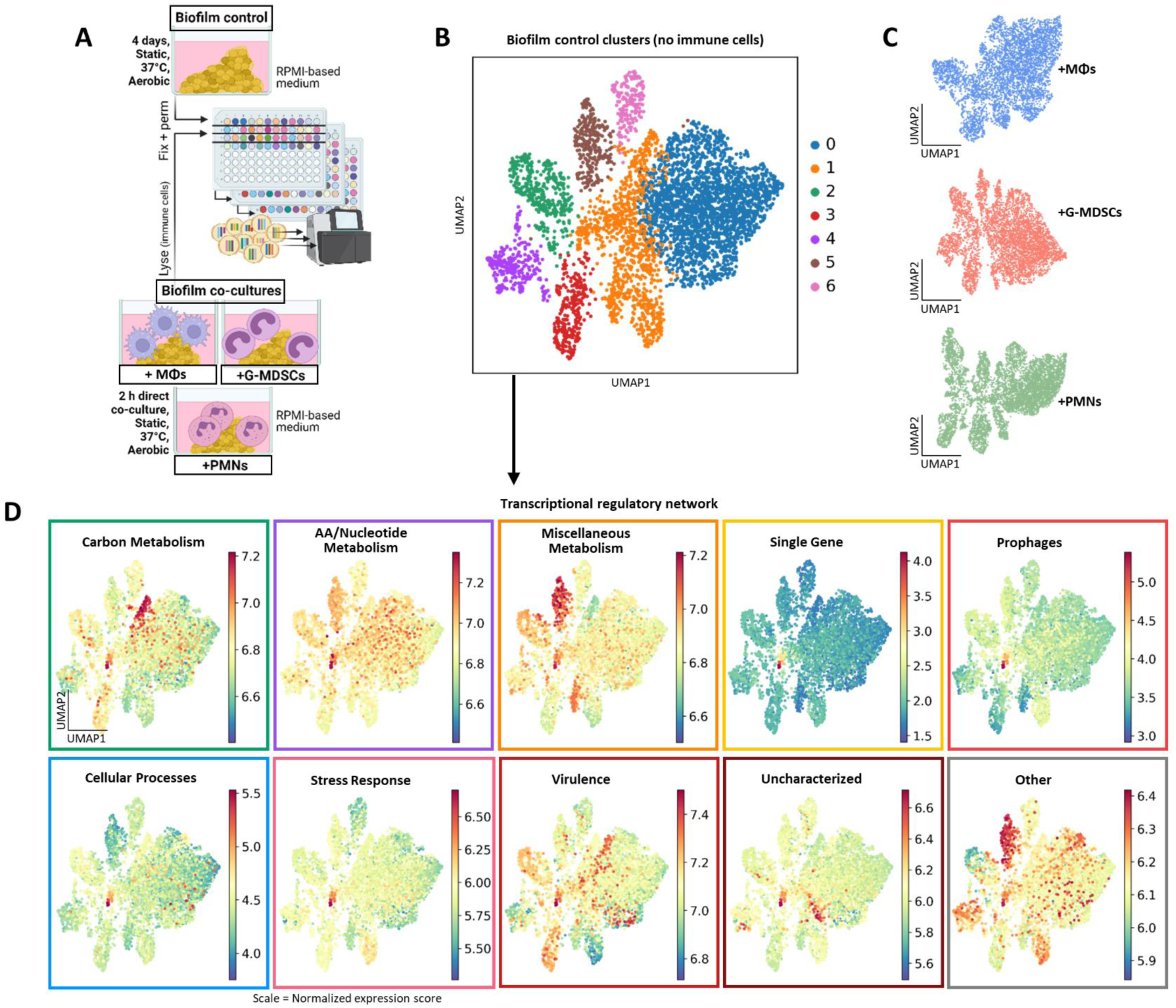
Biofilm shows coordinated transcriptional regulation. (A) *S. aureus* biofilm was grown for 4 days in RPMI-based medium under static, aerobic conditions at 37°C with daily medium replenishment. After 4 days, biofilm was directly co-cultured with 5×10^5^ mouse bone marrow-derived macrophages (MΦs), granulocytic myeloid-derived suppressor cells (G-MDSCs), or neutrophils (PMNs) for 2 h, whereupon immune cells were lysed with water. Cells from biofilm control (no immune cells) and the co-cultures were fixed overnight before permeabilization under identical conditions. Cells from each respective sample were separate for the first round of barcoding, then combined for the second and third rounds. The combined samples were processed through to sequencing. (B) UMAP plot of the biofilm control, with colors denoting transcriptional subpopulations identified with the Leiden algorithm. (C) UMAP plots of independently clustered biofilms co-cultured with immune cells. Responses to each immune population are explored in more detail in Figure 6. (D) Transcriptional regulatory categories for the biofilm control are overlaid on UMAP plots. Categories correspond to the schematic in Figure 3A. Color represents normalized expression score per cell.

We first characterized the biofilm itself to identify differences between the various transcriptional clusters using iModulon analyses (Figure 4D). This revealed a complex picture of transcriptional regulation across biofilm subpopulations. Carbon metabolism and virulence regulatory networks showed strong coordination across the clusters, where expression scores were highest in clusters 2 and 3 while low in clusters 4, 5, and 6. Cluster 1 expressed a transcriptional signature that encompassed all of the regulatory categories, and a stress response appeared to be moderately uniform across most biofilm clusters with a hotspot in cluster 1. Cluster 5 showed a prominent signature that was classified as miscellaneous metabolism (Figure 4D). These initial high-level analyses help illustrate regulatory heterogeneity within biofilm, as a means to aid in understanding full transcriptional diversity.

### Biofilm subpopulations are characterized by diverse gene expression profiles

After obtaining an overview of biofilm transcriptional regulation with iModulon analyses, we next examined unique marker genes for each biofilm cluster to gain deeper insight into the functional state of each population, without influence from a planktonic comparison. Differential expression was performed for a given cluster vs. all others to identify marker genes (Figure 5A). Of note, some biofilm clusters contained a greater number of statistically significant genes than others, and not all sets of marker genes translated to a meaningful classification. This may be influenced by clustering parameters, which were carefully considered to optimize unambiguous identification of unique marker gene sets (see Methods and Figure S6). The top 5 genes for each biofilm cluster are listed in Table 1.

**Figure 5.**
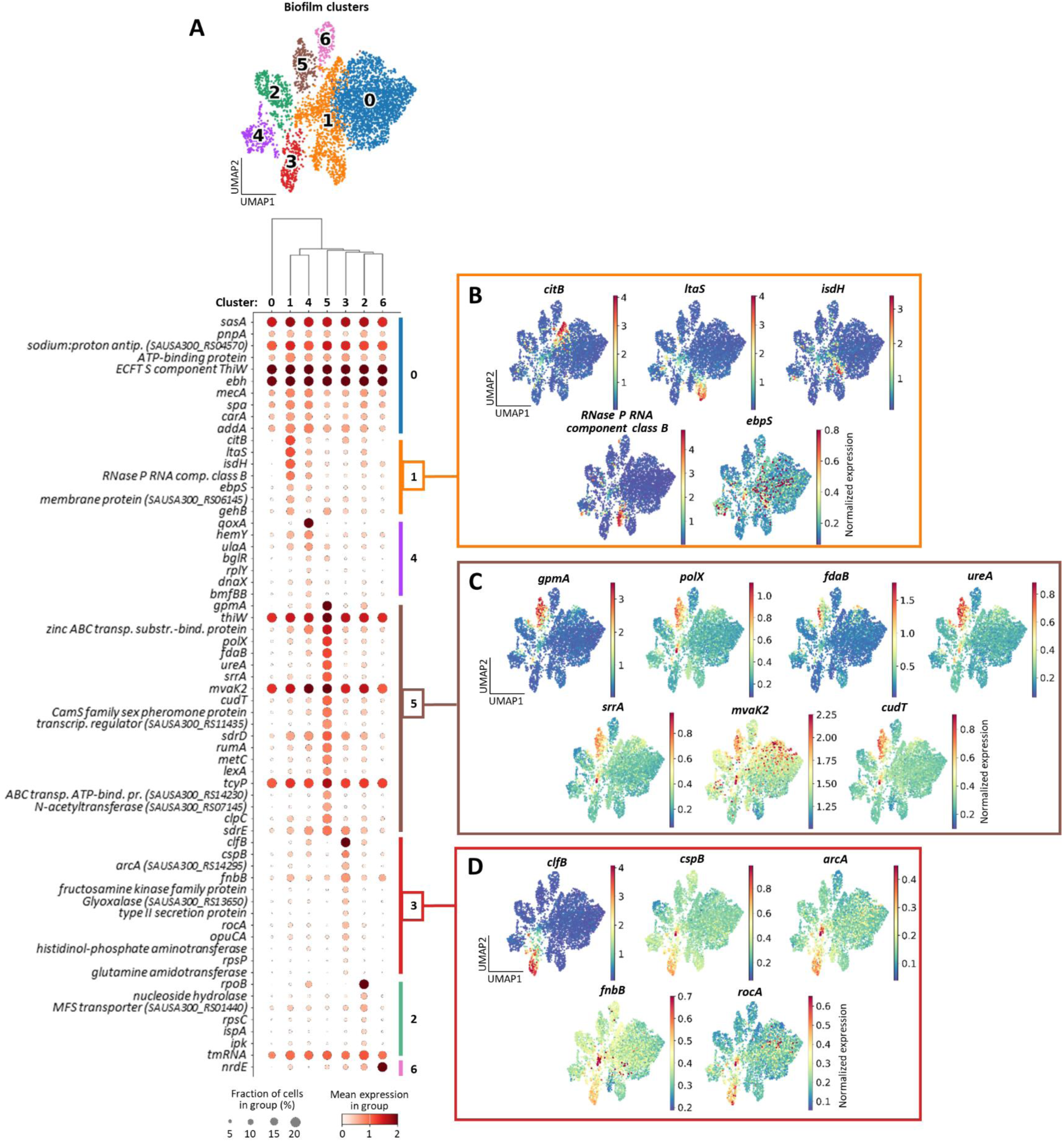
Biofilm subpopulations are characterized by diverse gene expression profiles. (A) Dot plot with top marker genes per cluster from the biofilm control sample. All marker genes have adjusted *p*-value ≤ 0.05. The UMAP plot is shown above for reference, and clusters are arranged in the plot according to transcriptional relationship, as illustrated in the dendrogram. A more detailed list of genes can be found in Table 1. ECFTS = energy coupling factor transporter S. (B, C, D) Marker gene expression overlaid on the UMAP plot for cluster 1 (B), cluster 5 (C), and cluster 3 (D). Color represents normalized expression per cell.

**Table 1.**
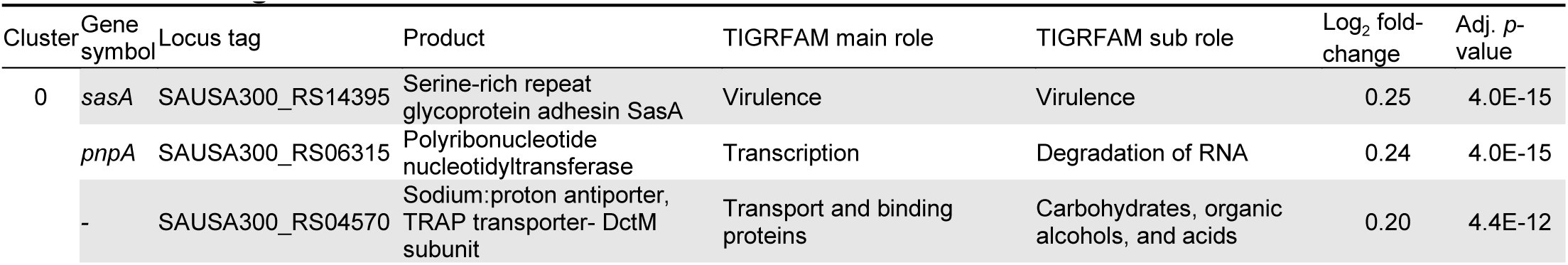
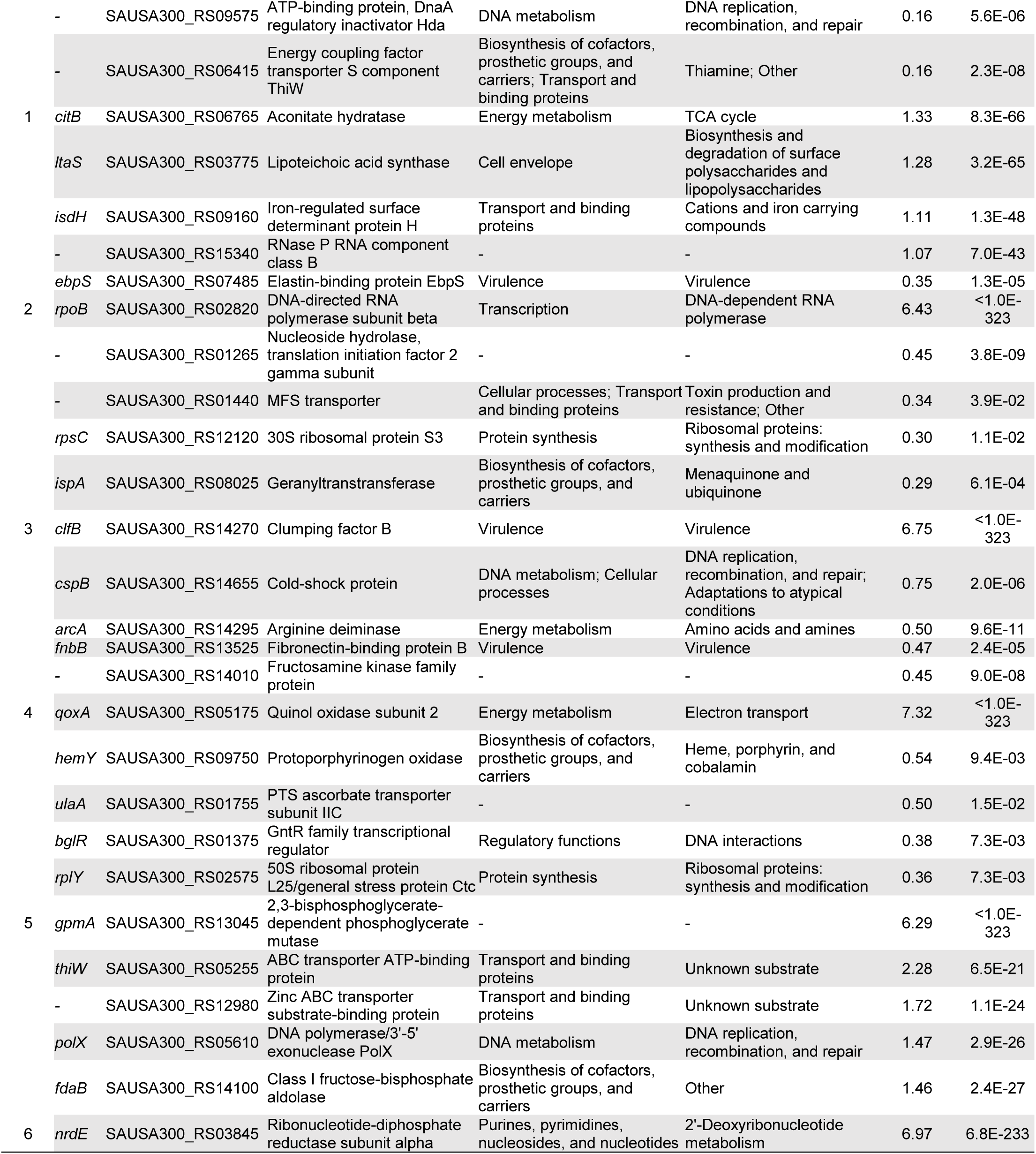
Marker genes from biofilm UMAP clusters.

Cluster 1 (Figure 5B) was identified as a transcriptionally active population, which was supported by ribonuclease P expression, an important enzyme in tRNA maturation.^122^ This cluster was also associated with *citB* expression, suggesting an active TCA cycle under catabolite control protein E (CcpE) regulation during low glucose conditions present in biofilm (Figure 2H).^123^ Other markers included lipoteichoic acid synthesis (*ltaS*), heme sequestration under iron limitation (*isdH*), and elastin binding/surface attachment (*ebpS*).^124–126^ Identifying cluster 1 as the most active population was corroborated by iModulon analyses (Figure 4D), which showed modest expression scores throughout all regulatory categories. Cluster 5 (Figure 5C) revealed a signature for alternative metabolism under stress. For example, *gpmA* and *fdaB* were enriched in cluster 5 and encode manganese-independent isozymes for two steps in glycolysis, suggesting metabolic activity under manganese limitation.^127,128^ Elevated expression of the DNA polymerase gene *polX* indicates cells undergoing replication or repair.^129^ Interestingly, PolX activity is manganese-dependent, suggesting that cells in cluster 5 could be prioritizing manganese usage under limiting conditions to maintain activity, although this remains speculative. Upregulation of the mevalonate pathway (*mvaK2*) further supports cellular activity in cluster 5.^130^ Besides an increase in genes involved in limiting nutrient stress, the expression of *ureA*, *srrA*, and *cudT* reflect responses to acidic, nitrosative and hypoxic, and osmotic stress, respectively,^131–133^ and were reflected in the iModulon network analyses (Figure 4D). Biofilm cluster 3 (Figure 5D) was enriched for genes involved in virulence and amino acid metabolism. For example, clumping factor B (*clfB*) and fibronectin-binding protein B (*fnbB*) are well-known virulence mechanisms of *S. aureus* involved in colonization and biofilm development.^92,93^ The *cspB* gene codes for a cold-shock protein with implications in small colony variant (SCV) formation that is a hallmark of biofilm cells.^134^ Several genes involved in amino acid metabolism were increased in cluster 3, including *arcA*, which converts arginine to citrulline producing ATP, CO_2_, and NH_3_, as well as *rocA* which generates glutamate from proline.^135,136^ Again, these classifications were supported by iModulon regulatory network analysis (Figure 4D). Overall, these complex gene expression patterns support a dynamic and heterogeneous transcriptional profile in biofilm.

### Biofilm differentially responds to distinct immune pressures

To explore how biofilm transcriptional profiles adapt to different immune pressures at single-cell resolution, we projected the bacterial cells from each biofilm co-culture condition (+MΦs, +G-MDSCs, and +PMNs) onto the biofilm control UMAP (Figures 4 and 5). This provided a controlled basis for comparison of each condition (Figure 6A). All biofilms co-cultured with immune cells consistently mapped to the control UMAP with nearly equivalent distributions of bacterial cells across the different clusters, validating no bias in the system (Figure 6B). As previously mentioned, biofilm cluster 1 contained the most transcriptionally active cells as depicted by overlaying total mRNA counts onto the UMAP space (Figure 6C, left). Trajectory analysis also revealed a pseudotime convergence to cluster 1 (Figure 6C, middle and Figure S4B) that corresponded with increased expression of ATP synthase genes (Figure 6C, right), which are important for influencing immune cell activation and biofilm persistence.^65,137^ Together, these findings suggest that the active biofilm population in cluster 1 experiences the most extensive transcriptional changes in response to immune pressure and was the focus of subsequent analysis. It is more challenging to compare less transcriptionally active clusters due to lower statistical power, which supports why cluster 1 was pursued.

**Figure 6.**
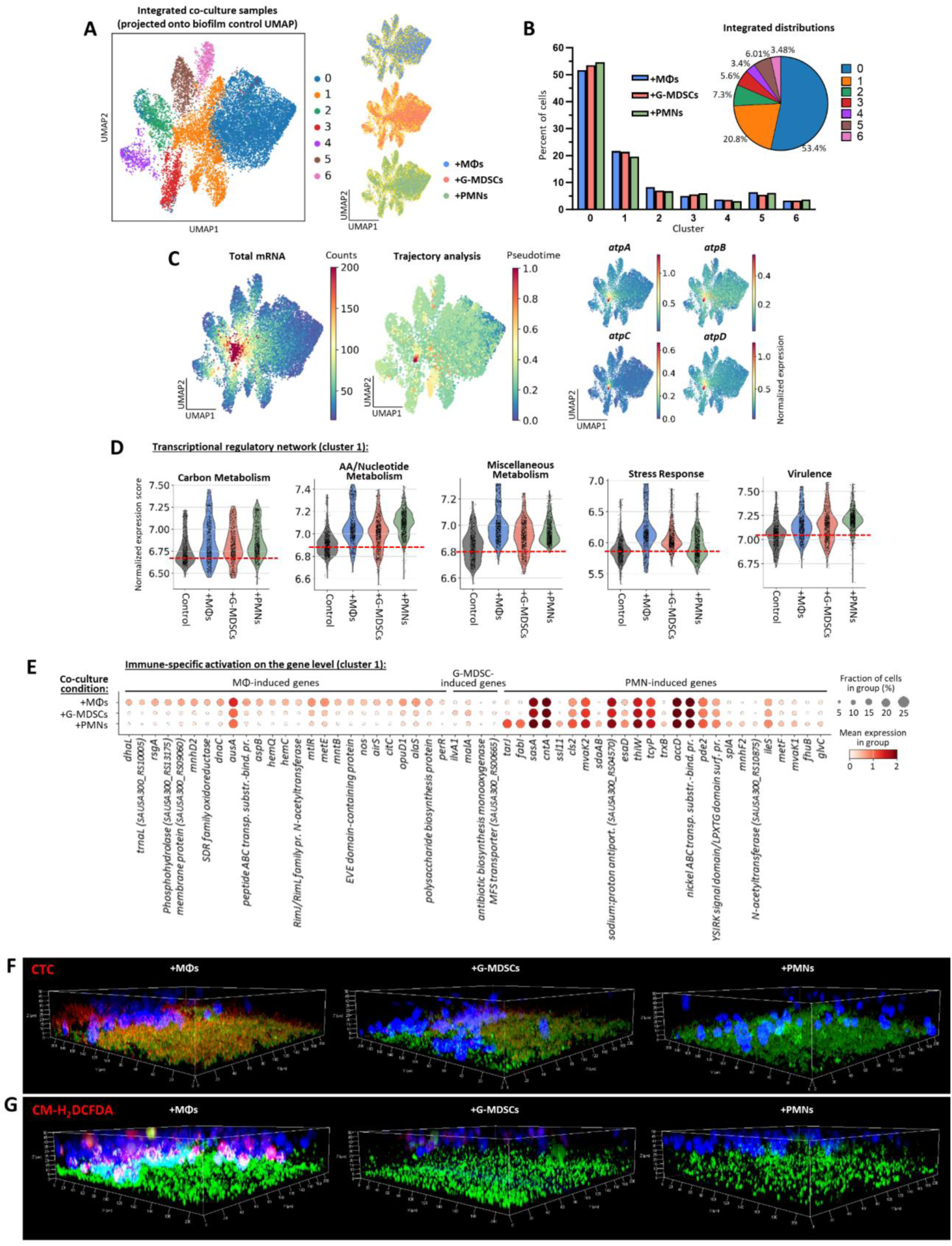
Biofilm differentially responds to distinct immune pressures. (A) Integrated UMAP plot of biofilms co-cultured with MΦs, G-MDSCs, and PMNs. The biofilm control UMAP (Figure 4B) was used as a template on which the co-culture samples were projected (using the *ingest* function of Scanpy). To the right, each individual co-culture condition is colored separately, with the other two co-culture conditions shown in yellow. (B) Bar plot showing the percentage of cells from each biofilm co-culture condition within each cluster. The pie chart shows combined cluster distributions for all co-culture samples. (C) (Left) Overlay of total mRNA counts on the integrated UMAP plot, with the highest number of captured transcripts present in cluster 1. (Middle) Trajectory analysis with the integrated biofilm-leukocyte co-cultured biofilm samples (Palantir algorithm), which identified a differentiation pathway that converges to cluster 1 upon immune cell exposure. (Right) Expression of *atpA/B/C/D* genes is concentrated in cluster 1, where the trajectory converges. (D) Transcriptional regulatory category expression within cluster 1 for the immune co-cultured biofilms compared to the biofilm control. Categories correspond to the schematic in Figure 3A. Red dashed lines depict the average of the biofilm control for reference. (E) Top genes activated in biofilm cluster 1 in response to leukocyte exposure. All genes have adjusted *p*-value ≤ 0.05 for differential expression (log_2_ fold-change) compared to the other co-culture conditions. (F-G) Evaluation of respiration and ROS activity using CTC (F) and CM-H_2_DCFDA (G) dyes in biofilm-leukocyte co-cultures with MΦs, G-MDSCs, or PMNs. Z-stack images were acquired (1 μm sections) and used to construct 3-D images. Color adjustments were applied uniformly across all images of the same experiment.

iModulon network activity across the different biofilm co-culture conditions revealed increased metabolic activity in cluster 1 in response to all three immune populations compared to the biofilm control, most prominently in utilization of miscellaneous amino acids and/or nucleotide sources (Figure 6D). In contrast, stress response and virulence pathways were divergently regulated in response to the three immune populations. Specifically, a stronger stress response in biofilm cluster 1 was observed following MΦ exposure, whereas PMNs induced a heightened virulence response (Figure 6D). G-MDSCs elicited the least perturbations in biofilm transcriptional profiles from the control. This was also confirmed by differential gene expression for each biofilm co-culture condition, where G-MDSCs induced minimal changes within biofilm (Figure 6E) consistent with the iModulon network analysis and the known ability of G-MDSCs to promote *S. aureus* biofilm persistence by their anti-inflammatory activity.^31,32^

MΦ co-culture elicited the most unique differentially expressed genes in biofilm compared to G-MDSCs and PMNs (Figure 6E). With regard to metabolism, MΦs upregulated *dhaL* that generates pyruvate from glycerol.^138^ MΦ co-culture also induced evidence of a stringent response with increased *rsgA* and *metE* expression, which encode a ribosome-associated GTPase that inhibits translation upon sensing (p)ppGpp and a methionine biosynthesis gene tied to stringent conditions, respectively.^139,140^ Genes for aspartate biosynthesis (*aspB*) and a non-ribosomal peptide synthase producing a protease inhibitor (*ausA*), both with ties to virulence, were also induced in biofilm specifically following MΦ exposure.^141,142^ However, the most upregulated genes during MΦ co-culture involved respiration and oxidative stress, where heme biosynthesis (*hemC/Q*) and nitric oxide synthase (*nos*) suggest active respiration under oxygen limiting conditions and oxidative stress elicited by MΦ activation.^143–145^ Genes involved in mannitol metabolism (*mtlR*) and manganese acquisition and competition (*mntB*) also imply osmotic and redox pressure, with *mntB* suggesting superoxide dismutase activation.^146,147^ Additional upregulated genes implicated in ROS detoxification include a regulator of staphyloxanthin production (*airS*), a regulator of peroxide resistance (*perR*), and a DNA helicase (*dnaC*) for replication and repair from oxidation-induced damage.^148–150^ Collectively, this suggests an adaptation to evade MΦ-mediated ROS production.

PMN co-culture also induced a unique transcriptional response in biofilm cluster 1 compared to MΦs and G-MDSCs (Figure 6E). Upregulation of several metabolic genes related to amino acid catabolism (*sdaAB*) and methionine (*metF*) were observed.^140,151^ Several respiration and oxidative stress genes were additionally increased, including staphylopine metal acquisition (*cntA*) and thioredoxin reductase (*trxB*).^152,153^ However, the most prominent genes upregulated during PMN co-culture are involved in cell wall maintenance and virulence, including *tarJ*, responsible for the rate-limiting step in CDP-ribitol synthesis for wall teichoic acids and *fabI*, a critical rate-limiting enoyl-ACP reductase for fatty acid synthesis.^154–156^ Interestingly, the activities of both *tarJ* and *fabI* require the oxidation of NADPH to NADP^+^, indicating important regulation of cell redox state. Additional upregulated genes involved in cell wall maintenance include mevalonate synthesis (*mvak1/2)*, which affects both cell wall synthesis and membrane stabilization, and cardiolipin synthase (*csl2)*.^157,158^ Increased cardiolipin synthase activity has been shown to inhibit PMN chemotaxis by reducing phosphatidylglycerol on the bacterial membrane, which is a chemoattractant.^159^ Virulence genes induced in biofilm cluster 1 following PMN co-culture include a cell surface protein involved in surface attachment (*sasA*), a serine protease (*splA*), and superantigen-like protein 11 (*ssl11*).^106,160,161^ Similar to cardiolipin induction, SSL11 has been shown to arrest PMN motility by inducing adhesion without oxidative burst.^160^

Biofilm cluster 0 contained roughly 50% of the cells in the dataset. Transcriptional network analysis and quantification of total mRNA transcripts from cells in cluster 0 indicated that this was a metabolically and transcriptionally dormant population. Further studies into this group of cells could address the controversial issue of what defines a persister cell.^162,163^ While these cells were not dead since the RNA would have degraded, their overall low activity suggests at least a portion may be persisters. Top marker genes for this cluster (Figure 5A) included several surface proteins with known roles in adherence and virulence (*sasA*, *ebh*) and an exoribonuclease for RNA degradation (*pnpA*). Interestingly, cluster 0 exhibited evidence of ‘reawakening’ following immune cell exposure (Figure S7) As evident by a conserved upregulation of a RNA polymerase component previously shown to be correlated with a planktonic growth trajectory (*rpoB*, Figure 3F), potentially suggesting reanimation to a more metabolically active population although this remains speculative.

The application of BaSSSh-seq to *S. aureus* biofilm co-cultures revealed the ability of biofilm to adapt and uniquely respond to distinct immune populations. Whereas anti-inflammatory G-MDSCs elicited minimal transcriptional changes, MΦs induced a prominent stress response to regulate respiration and oxidative damage, and PMNs induced genes related to cell wall maintenance and virulence. These observations were validated by confocal microscopy where *S. aureus* biofilms directly co-cultured with MΦs, G-MDSCs, or PMNs were stained with CTC (5-cyano-2,3-ditolyl tetrazolium chloride) or CM-H_2_DCFDA (6-chloromethyl-2’,7’-dichlorodihydrofluorescein diacetate) to broadly probe respiration and ROS, respectively. Specifically, biofilm co-culture with MΦs led to increased respiration within biofilm (Figure 6F) and the largest ROS signature (Figure 6G) compared to G-MDSCs and PMNs, consistent with our transcriptomic analyses. Collectively, these findings demonstrate the sensitivity and selectivity of BaSSSh-seq to identify unique transcriptional alterations in *S. aureus* biofilm that can be used to examine how biofilm adapts transcriptional profiles in response to immune pressure.

## DISCUSSION

Here we present BaSSSh-seq, a bacterial scRNA-seq method incorporating a plate-based barcoding system with rRNA depletion. BaSSSh-seq was applied to study *S. aureus* biofilm heterogeneity and immune interactions, an advance from previous demonstrations of bacterial scRNA-seq on planktonic cells. In this application, we captured vast transcriptional heterogeneity within biofilm compared to planktonic growth and showed the ability to detect distinct biofilm responses tailored to different immune cell populations. In addition to the technical advances in scRNA-seq methodology, our analyses present a conceptual advance towards the understanding of complex biofilm communities through incorporation of new computational pipelines that enable high-level regulatory network visualization and trajectory inference paired with gene-level expression quantification.

Our BaSSSh-seq methodology was validated by literature comparing alterations in gene expression and metabolism during biofilm vs. planktonic growth. Moreover, our subsequent analyses laid the groundwork for exploration beyond simple validation. A current lack of understanding exists surrounding the intricately coordinated cellular networks that govern biofilm growth, stemming from inadequate high-throughput methods to measure the stochastic interactions between discrete subpopulations. A promising avenue for insights lies in the coupling of bacterial scRNA-seq with transcriptional regulation analysis, as implemented in our study. The iModulon-based assessments enable cross-population relationships to be quantified and visualized. Furthermore, trajectory analysis provides another means to understand signaling dynamics, especially when linked to gene expression. While only a subset of genes correlating with the trajectory were discussed, many more remain unexplored (Tables S1-S2). Several of these genes encode uncharacterized proteins that could potentially play key roles in biofilm formation and may represent attractive anti-biofilm therapeutic or prophylactic targets. An important future direction towards a better understanding of biofilm growth is to perform BaSSSh-seq during different stages of biofilm growth to assess temporal alternations in gene expression, the transcriptional regulatory network through iModulons, and clustering patterns during maturation. We did not detect many genes previously identified to be important during biofilm formation, such as the *icaABCD* and *cidAB* operons, which is likely because established biofilms were examined in this study.^54,57^ Relating transcriptionally defined clusters to spatially defined microstructures and regions throughout the various stages of biofilm development would augment our understanding of biofilm growth and signaling, which could be achieved by constructing fluorescent reporters for genes that are enriched in distinct clusters.

BaSSSh-seq successfully generated powerful visualizations of biofilm transcriptional regulation paired with gene-level analyses for subpopulation characterization. The heterogeneity and coordinated patterns of gene regulation observed across biofilm clusters overwhelmingly illustrate how the ensemble-averaged expression from traditional bulk RNA-seq is insufficient. Accordingly, single-cell resolution also provides quantitative information on relative population sizes, a metric that is lost in bulk methods. Although many biofilm cells displayed a transcriptionally dormant phenotype (cluster 0), we focused our efforts on more active biofilm populations and how they interacted with the immune response. Our analyses demonstrated that biofilm undergoes dramatic transcriptional alterations that are tailored to the immune cell encountered. Although speculative, it is intriguing to consider that the most metabolically active biofilm clusters were responsive to macrophage and PMN challenge, since these immune populations are major producers of ROS, RNS, and proteases that place strong pressures on bacteria.^119,144,148,149^ In contrast, G-MDSCs do not exhibit antibacterial activity, so the biofilm does not need to expend substantial energetic resources to transcriptionally respond to this non-threat.^31,32^ These findings have significant potential to inform more effective immunomodulatory therapies and support the concept of nutritional immunity described in the literature.^164^ Future efforts will move *in vivo*, to explore the diversity of *S. aureus* adaptation and immune responses across different tissue niches.

Although highly functional, areas for improvement remain throughout the BaSSSh-seq methodology and analyses. For example, the number of barcoded cells with appreciable numbers of mRNA reads in biofilm samples was low. Insights from our comparisons of biofilm and planktonic cultures suggest this results from decreased transcriptional activity within biofilm. Nonetheless, membrane permeabilization conditions prior to barcoding could be more thoroughly studied to improve time and temperature for maximal barcode diffusion and RNA capture. Additionally, the barcoding could be expanded to 384-well plates to increase cell capacity by >60X. Sequencing depth also impacts the capture and detection of low-level transcripts, and with incorporation of rRNA depletion we improved cost efficiency and information content for sequencing runs, permitting usage of a mid-output kit on an Illumina NextSeq 500/550 series platform. However, availability of larger sequencers and kits exist for increasing sequencing depth >200X. As discussed further in the Methods, an inherent background noise exists, evident in the UMAP overlays in Figures 2-5 where many genes were expressed at baseline levels throughout all clusters. This limitation restricted the statistical power of some analyses, and improvements would allow for higher confidence in identifying targets for experimental validation. Reduction in noise levels could be realized through adjustment of randomer concentrations in both reverse transcription and second strand synthesis steps, fragmentation conditions used in library prep, and/or modification of alignment parameters. Clustering itself could be further optimized to identify more meaningful classifications through further adjustments to parameter settings and/or future advances in clustering tools and algorithms. From a technical perspective, exploration of long-read sequencing presents a promising avenue that would allow fragmentation to be bypassed, leading to substantial noise reduction while potentially providing new insights into large-scale operon architecture. Several limitations are also evident from an experimental standpoint. First, as noted above, this study examined mature biofilm to assess how various immune cell subsets altered transcriptional programs. Performing BaSSSh-seq at regular intervals during biofilm development could provide new insights into fundamental populations that expand at key steps (i.e., attachment, exodus, and expansion). Second, spatial information about how specific biofilm transcriptional clusters relate to structural attributes (i.e., attachment, tower formation) is an interesting area to pursue as the resolution of spatial transcriptomic approaches improve. Based on the nature of this work describing BaSSSh-seq as a resource, the importance of specific *S. aureus* genes in biofilm biology or metabolism were not assessed, although we did validate changes in biofilm metabolism, respiration, and ROS as an initial step. Finally, only one co-culture interval of biofilm and immune cells was examined (2 h) as a proof-of-concept for biofilm adaptation; however, the kinetics of these changes could be explored in future studies.

Overall, the BaSSSh-seq method coupled with powerful computational approaches facilitates the high-throughput study of biofilm transcriptional heterogeneity at a new resolution. The datasets provide a rich resource for the biofilm community to explore, and the optimized protocols and analyses provide a mechanism to aid in identification of new therapeutic targets and strategies.

## ACKNOWLEDGMENTS

This work was supported by the National Institutes of Health/National Institute of Neurological Disorders and Stroke F32NS126302 to LEK and the National Institute of Allergy and Infectious Diseases 3P01AI083211 (Project 4) to TK. The University of Nebraska DNA Sequencing Core receives partial support from the National Institute for General Medical Science INBRE - P20GM103427-19 grant as well as The Fred & Pamela Buffett Cancer Center Support Grant - P30CA036727. We thank Drs. Paul D. Fey and Vinai C. Thomas for their expert discussions and suggestions on data analysis and interpretation. We thank Jennifer Endres for her support with equipment, reagents, and database curation. We acknowledge Dr. Eric Tom and Nichole Brandquist for their technical assistance on sequencing library QC and confocal imaging, respectively. We acknowledge Dr. Anna Kuchina and Sydney Blattman for sharing their expertise during initial experimental setup. We appreciate the discussions and suggestions from current and former members of the Kielian Laboratory and others at UNMC including Drs. Blake Bertrand, Christopher Horn, Gunjan Kak, Prabhakar Arumugam, as well as Zachary Van Roy and Cleofes Sarmiento.

## AUTHOR CONTRIBUTIONS

LEK and TK designed the study. LEK conducted experiments, performed data analysis, and wrote the manuscript. Both authors edited and approved the final manuscript.

## METHODS

### RESOURCES

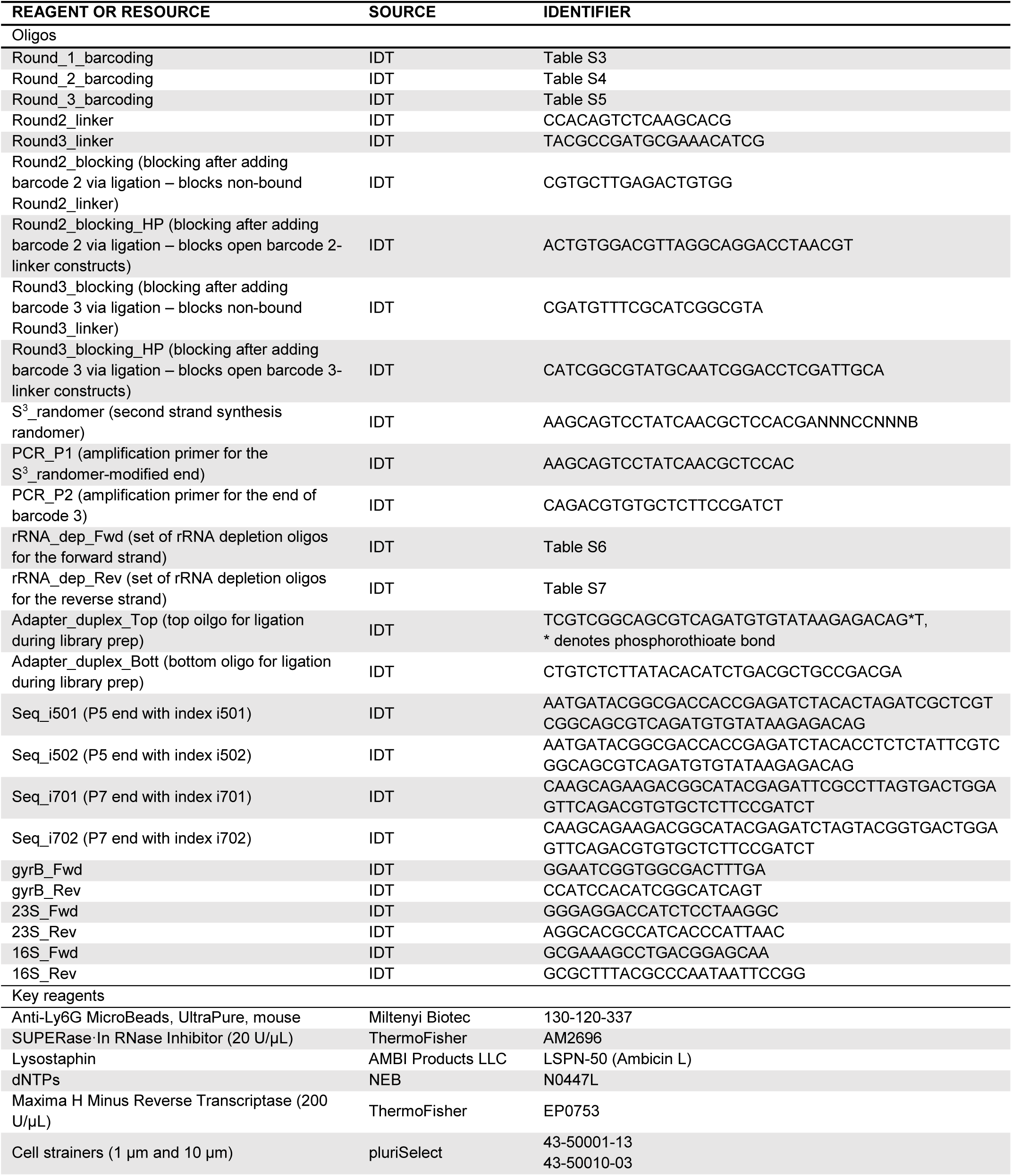
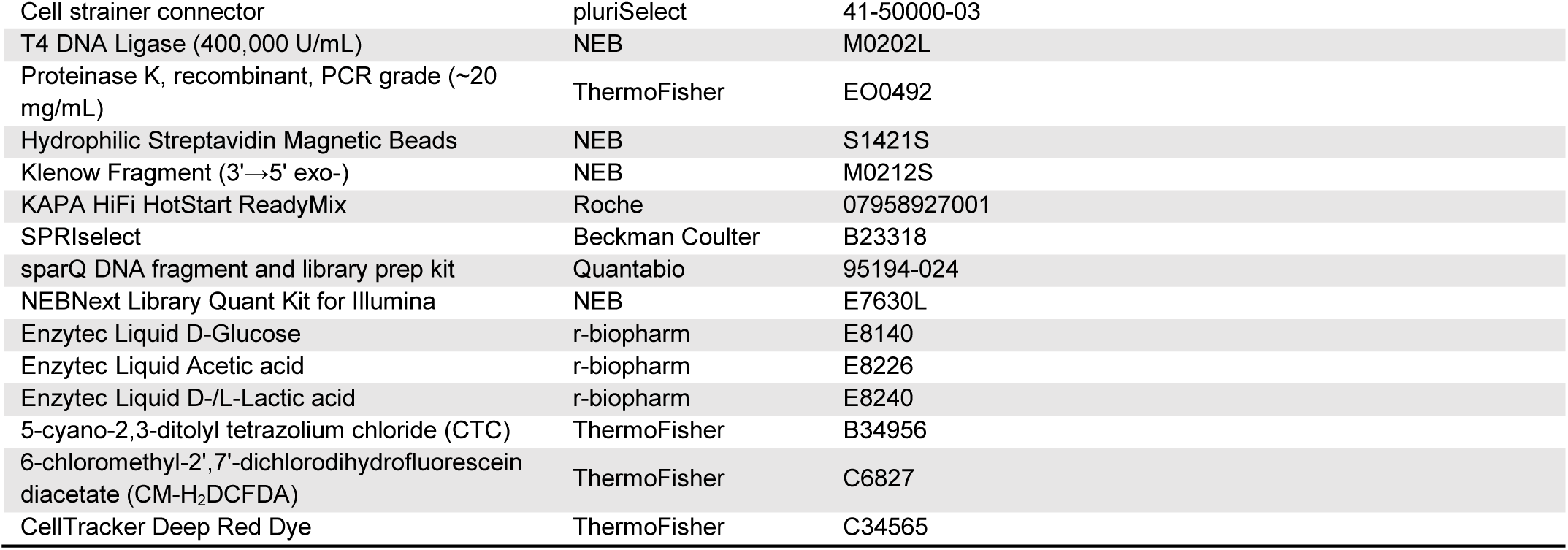

### DATA AVAILABILITY

Sequencing data and processed count matrices will be made public upon publication.

### EXPERIMENTAL DETAILS

#### Bacterial strains

All sequencing experiments were performed with *S. aureus* USA300 LAC-13C.^165^ For confocal microscopy, *S. aureus* GFP pCM29 and dsRed pVT1 expressing strains were used as previously described, with plasmids maintained during *in vitro* growth with 10 ug/mL chloramphenicol.^166,167^

#### RPMI-based medium

The RPMI-based medium used throughout all experiments for biofilm and planktonic cultures was RPMI-1640 supplemented with 10% heat-inactivated FBS, 1% L-glutamate, and 1% HEPES.

#### *In vitro* biofilm growth

24-well plates were coated overnight in 20% human plasma in 10X PBS at 4°C. Plasma coating solution was removed prior to seeding each well with 600 μL of *S. aureus* from an overnight culture grown for 16-18 h at 37°C under aerobic conditions at 250 rpm using a 1:10 volume:flask ratio (25 mL RPMI-based medium in a 250 mL baffled flask) diluted 100X in RPMI-based medium. Plates were incubated under static, aerobic conditions at 37°C for 4 days. Each day, 270 μL of spent medium was carefully removed from each well without disturbing biofilms, whereupon 300 μL fresh medium was carefully added to avoid disturbing the biofilm. For confocal microscopy experiments, biofilms were grown as described in 8-well chamber slides with 400 μL total liquid volume.

#### Planktonic growth

A single colony of *S. aureus* was inoculated into 25 mL RPMI-based medium at a 1:10 volume:flask ratio (250 mL baffled flask) for overnight (16-18 h) aerobic growth at 37°C and shaking at 250 rpm. The following day, 250 μL of this overnight culture was inoculated into 25 mL fresh RPMI-based medium for outgrowth to exponential phase (3-3.5 h, OD_600_=0.35) under the same conditions as the overnight culture.

#### Overview of the BaSSSh-seq protocol for comparing biofilm and planktonic growth

Cells from biofilm and planktonic samples were collected by brief centrifugation at 12,000x*g* for 1 min, and immediately resuspended in fixation buffer (4% formaldehyde in 1X PBS) for overnight incubation at 4°C before permeabilization the next morning (both fixation and permeabilization performed in parallel and under identical conditions). Cells from biofilm and planktonic cultures were kept separate for the first round of barcoding (each sample to 24 wells of the first barcoding plate), then combined for the second and third rounds. The combined samples were processed through second strand synthesis, rRNA depletion, and library prep to sequencing. In total, ∼200,000 cells were processed for sequencing.

#### Isolation of primary MΦs, G-MDSCs, and PMNs

All immune cell types were isolated from the bone marrow of 8-10 wk old WT C57BL/6J mice (RRID:IMSR_JAX:000664) as previously described.^168^ For MΦs, bone marrow cells were incubated in RPMI-based medium supplemented with 1X antibiotic/antimycotic solution (100 U/mL penicillin G, 100 μg/mL streptomycin, 0.25 μg/mL amphotericin B), 50 μM 2-mercaptoethanol, and M-CSF from L929 cells for 7 days at 37°C and 5% CO_2_, with medium changes on days 3 and 5. For G-MDSCs, bone marrow cells were incubated in RPMI-based medium supplemented with 1X antibiotic/antimycotic solution, 50 μM 2-mercaptoethanol, and 40 ng/ml each of G-CSF and GM-CSF for 4 days at 37°C and 5% CO_2_, with 40 ng/ml of IL-6 added on day 3. After 4 days, G-MDSCs were purified with Anti-Ly6G MicroBeads. For PMNs, bone marrow was isolated and immediately purified with Anti-Ly6G MicroBeads.

#### Overview of the BaSSSh-seq protocol for biofilm-leukocyte co-cultures

*S. aureus* biofilm was directly co-cultured with 5×10^5^ primary MΦs, G-MDSCs, or PMNs for 2 h. After co-culture, all cells (bacteria and immune) were collected and briefly centrifuged at 12,000x*g* for 1 min. The cell mixture was resuspended in water for 10 min with brief, intermittent vortexing to preferentially lyse immune cells to prevent eukaryotic RNA contamination. After another centrifugation at 12,000x*g* for 1 min, bacterial cells were immediately resuspended in fixation buffer (4% formaldehyde in 1X PBS) for overnight incubation at 4°C before permeabilization the next morning. Cells from the biofilm control (no immune cells) and co-cultures were fixed and permeabilized in parallel under identical conditions. Cells from each respective sample were separate for the first round of barcoding (each sample to 12 wells of the first barcoding plate), then combined for the second and third rounds. The combined samples were processed through second strand synthesis, rRNA depletion, and library prep to sequencing. In total, ∼400,000 cells were processed for sequencing.

#### Confocal microscopy

Biofilms were grown in 8-well glass chamber slides and were visualized during immune cell co-cultures using confocal laser scanning microscopy (Zeiss 710) with a 40X oil lens. Z-stack images were acquired (1 μm sections) and used to construct 3-D images. For 5-cyano-2,3-ditolyl tetrazolium chloride (CTC, respiration marker) staining, GFP-expressing bacteria were used for biofilm and immune cells were labeled with CellTracker Deep Red. After 4 days of biofilm growth, 180 μL of medium was carefully removed from biofilms, whereupon 100 μL of 4 mM CTC (final working concentration of 1 mM) followed by 100 μL of each leukocyte population (1.5×10^6^ cells) was carefully added for a final volume of 400 μL, and images were acquired within 10 min to prevent signal saturation. For 6-chloromethyl-2’,7’-dichlorodihydrofluorescein diacetate (CM-H_2_DCFDA, ROS marker) staining, dsRed-expressing bacteria were used for biofilm and immune cells were labeled with CellTracker Deep Red. After 4 days of biofilm growth, 180 μL of medium was carefully removed from biofilms, whereupon 100 μL of 40 μM CM-H_2_DCFDA (final working concentration of 10 μM) followed by 100 μL of each leukocyte population (1.5×10^6^ cells) was carefully added for a final volume of 400 μL, and images were acquired within 40 min to prevent signal saturation.

#### Metabolite measurements

Supernatants were collected from biofilm during initial inoculation and daily (prior to medium replenishment) for 4 days. Supernatants were collected from planktonic cultures during initial inoculation and every 30 min up to 6 h. Enzytec UV assay kits for Liquid D-Glucose (E8140), Liquid Acetic acid (E8226), and Liquid D-/L-Lactic acid (E8240) were used for quantification.

#### Solutions used throughout BaSSSh-seq processing from barcoding to sequencing

**PBS+RI:** 0.1 U/μL RI (SUPERase·In RNase Inhibitor) in 1X PBS

**Tris-HCl+RI:** 100 mM Tris-HCl pH 8.0 (same pH used throughout), 0.1 U/μL RI

**Permeabilization mix:** 100 mM Tris-HCl, 0.05 M EDTA, 0.25 U/μL RI, 40 ug/mL lysostaphin

**2X RT mix (600 μL):** 30 μL water (molecular biology grade, used throughout), 240 μL 5X RT buffer (provided with Maxima H Minus Reverse Transcriptase), 30 μL RI, 60 μL dNTPs, 180 μL 50% PEG8000, 60 μL Maxima H Minus Reverse Transcriptase

**Ligation mix (1.02 mL):** 295 μL water, 250 μL 10X T4 DNA Ligase buffer (provided with T4 DNA Ligase), 75 μL T4 DNA Ligase, 25 μL RI, 375 μL 50% PEG8000

**Wash buffer:** 0.1% Triton X-100 and 0.05 U/μL RI in 1X PBS

**2X Lysis buffer:** 20 mM Tris-HCl, 400 mM NaCl, 100 mM EDTA, 4.4% SDS

**2X BW buffer:** 10 mM Tris-HCl, 2 M NaCl, 1 mM EDTA, 0.1% Tween-20

**S^3^TE-TW buffer:** 10 mM Tris-HCl, 0.01% Tween-20, 1 mM EDTA

**S^3^ mix (440 μL):** 111.1 μL water, 88 μL 5X RT buffer (provided with Maxima H Minus Reverse Transcriptase), 176 μL 30% PEG8000, 44 μL 10 mM dNTPs, 4.4 μL 1 mM S^3^_randomer, 16.5 μL Klenow Fragment (3’→5’ exo-)

**PCR mix (440 μL):** 184.4 μL water, 17.6 μL 10 μM PCR_P1, 17.6 μL 10μM PCR_P2, 220 μL 2X KAPA HiFi HotStart ReadyMix

**T.1E:** 10 mM Tris-HCl, 0.1 mM EDTA

#### Barcoding

##### Cell fixation, permeabilization, and counting

Fixation was achieved using 4% formaldehyde in 1X PBS overnight at 4°C. The next morning, cells were briefly vortexed and centrifuged at 7,000x*g* for 5 min at 4°C (standard centrifugation conditions used throughout the entire barcoding process) and resuspended in 1 mL cold Tris-HCl+RI. Cells were centrifuged and washed again in Tris-HCl+RI. Next, cells were resuspended in 500 μL 0.04% Tween-20 in 1X PBS and incubated on ice for 3 min. A 1 mL volume of cold PBS+RI was added before centrifuging cells and resuspending in 300 μL permeabilization mix. Cells were held in permeabilization mix at 37°C for 15 min, with intermittent mixing. After permeabilization, 1 mL of cold PBS+RI was added before centrifugation. Cells were resuspended in 500 μL cold PBS+RI and 1 μL of 10% Tween-20 was added before another centrifugation and resuspension in 500 μL cold PBS+RI. Cells were stored on ice while counting was performed on a hemocytometer. Cells were diluted to ∼3×10^6^ per mL for barcoding.

##### Reducing cell clumping and aggregates during barcoding

Before each barcoding step and after ligation of the final barcode, bacterial cells were vortexed for 1 min and filtered through consecutive 10 μm and 1 μm cell strainers with gentle vacuum. Immediately prior to loading cells into the barcoding plates, cells were again vortexed for 1 min and briefly sonicated for 1-2 sec.

##### In-cell reverse transcription (barcode 1)

Barcode 1 was added during the initial in-cell capture of RNA transcripts through random hexamer-primed reverse transcription. Wells of a 96-well plate were loaded with 6 μL of 25 μM respective round_1_barcoding oligo, 10 uL of 2X RT mix, and 4 uL of cells (at ∼3×10^6^ per mL after vortexing, filtering, and sonicating). For experiments with multiple samples, the cells from distinct samples were kept separate for the first barcoding round for proper identification based on barcode 1 after sequencing. Several important factors influenced the selection of cell numbers for barcoding. The number of cells should not dramatically exceed the number of possible barcode combinations, or duplicated barcodes may arise in the sequencing data. However, in addition to unavoidable cell loss throughout the barcoding process, a large number of sequenced cells (more than half) contained an inadequate number of RNA reads and were filtered out of the analysis (Figure S3A and Figure S5A). This likely resulted from multiple factors including low transcriptional activity of cells in biofilm, incomplete permeabilization, and poor capture efficiency. The reverse transcription plate was incubated for 10 min at 23°C and 50 min at 50°C in a thermocycler (lid temperature set at 50°C). Following reverse transcription, cells from all wells were collected and pooled, and 9.6 μL of 10% Triton X-100 was added prior to centrifugation. Cells were resuspended in 1 mL cold PBS+RI and vortexed, filtered, and sonicated as described above.

##### In-cell ligations (barcodes 2 and 3)

Barcodes 2 and 3 were added by in-cell ligation with the prior barcode. For the first ligation reaction, 2.5 μL of 30 μM respective round_2_barcoding oligo and 2.5 uL of 30 μM round2_linker oligo were pre-annealed in each well of a 96-well plate by heating to 95°C for 2 min and cooling to 20°C at a rate of −0.1°C/sec in a thermocycler (standard conditions for each pre-annealing step). The 1 mL of pooled cells from reverse transcription was mixed with 1.02 mL ligation mix (after vortexing, filtering, and sonicating), and 20 μL was added to each well containing the 5 μL pre-annealed barcode-linker mix. The plate was incubated at 37°C with shaking for 30 min. Then, 5 μL of blocking mix (40 μM round2_blocking and 40 μM pre-annealed round2_blocking_HP in 2.5X T4 DNA Ligase buffer) was added to each well and incubated at 37°C with shaking for 30 min. Cells from all wells were then pooled and vortexed, filtered, and sonicated as described above. After the final vortex and sonication, 50 μL of fresh T4 DNA Ligase was added to the cell pool. For the second ligation reaction, 3 μL of 30 μM respective round_3_barcoding oligo and 3 μL of 30 μM round3_linker oligo were pre-annealed in each well of a 96-well plate. The pooled cells from the first ligation reaction were then aliquoted at 24 μL per well. The plate was again incubated at 37°C with shaking for 30 min. Then, 5 μL of blocking mix (45 μM round3_blocking and 45 μM pre-annealed round3_blocking_HP in 150 mM EDTA) was added to each well and incubated at 37°C with shaking for 30 min. Cells from all wells were then pooled and vortexed, filtered, and sonicated as described above.

##### Final washing and cell library construction

After vortexing, filtering, and sonicating the final barcoded cell mixture, 36 μL of 10% Triton X-100 was added before centrifugation. The supernatant was removed until ∼30 μL remained, to avoid aspiration of the cell pellet that was small and fragile at this step. The cell pellet was resuspended in 1.5 mL wash buffer and washing was repeated once to ensure adequate cleaning of the cells and removal of excess reagents. Next, the final cell pellet was resuspended in ∼200 μL PBS+RI, and cells were counted on a hemocytometer. Aliquots of 15,000-25,000 cells in 0.5 mL tubes were then brought up to 50 μL with 1X PBS and stored at −80°C until lysis. The number of cells per library was dependent on the experiment and how many cells were targeted for sequencing. As previously mentioned, a large portion of cells were filtered out based on low numbers of RNA reads per cell. After several sequencing runs and observing the numbers of cells filtered, cell library sizes were adjusted accordingly. Compiling cell libraries with a large number of cells is ideal for conserving reagents and costs; however, we observed that if cell libraries exceeded 25,000 cells, the downstream second strand synthesis step was error prone (Figure S1B-C). We believe this was caused by excess round_3_barcoding oligos that are free in solution, or not bound to captured RNA transcripts, but are still pulled out of cell lysates by their biotin tag and carried through second strand synthesis by the streptavidin beads.

##### Cell lysis

Cell libraries were flash thawed from −80°C, and 50 μL 2X lysis buffer and 10 μL proteinase K solution was added. Lysis reactions were incubated at 55°C for 2 h with shaking at 600-750 rpm. Lysates were directly stored at −80°C until purification and second strand synthesis.

#### Second strand synthesis

##### Purification of captured transcripts

The biotin tag on round_3_barcoding oligos facilitated transcript purification from the cell library lysates. 100 μL of Hydrophilic Streptavidin Magnetic Beads per library were aliquoted into separate 1.5 mL tubes for washing. Washing consisted of placing tubes on a magnetic stand for ∼1 min until beads were sufficiently pulled out of solution, removing the liquid, and resuspending beads in ∼360 μL of a particular wash buffer (note that different wash buffers were used throughout second strand synthesis). Washing was performed three times in 1X BW buffer before final resuspension of beads in 100 μL of 2X BW buffer. 2 μL of RI was added to the final suspension. Cell library lysates were briefly thawed in a 37°C bath until the solution became clear (the solution was initially turbid and white when cool). Then, 5 μL of 10 mM PMSF was added to each lysate library and incubated at room temperature for 10 min to inhibit proteinase K activity. After 10 min, 100 μL of washed bead suspension was added to each lysate library in 0.5 mL tubes and biotin-streptavidin binding occurred during shaking at ∼750 rpm at room temperature for 1 h. Beads tended to settle over time, so tubes were manually inverted every ∼10 min to mix. After the 1 h binding, beads were washed in 1X BW buffer twice followed by S^3^TE-TW buffer twice, with 5 min shaking at ∼750 rpm after resuspension during each wash.

##### Denaturing and second strand synthesis

Reverse transcription in the first barcoding round resulted in a hybrid RNA-cDNA construct. For second strand synthesis to occur, the captured RNA from the RNA-cDNA constructs was denatured in order for random priming of the cDNA (with oligo S^3^_randomer) to carry out second strand synthesis, as originally described.^29^ After the final wash in S^3^TE-TW buffer in the previous step, beads with captured transcripts were resuspended in 500 μL 0.1 M NaOH (prepared fresh on the day of use) and incubated at room temperature for 5 min with shaking at ∼750 rpm with inversion every 1 min. Beads were then washed twice in S^3^TE-TW buffer and twice in 10 mM Tris-HCl pH 8.0. After the last wash, beads were resuspended in 220 μL S^3^ mix per library and incubated at 37°C for 1 h with shaking at 600-700 rpm, and tubes were inverted to prevent bead settling every ∼10 min. After second strand synthesis, beads were washed twice in S^3^TE-TW buffer and once in water (molecular biology grade, used throughout).

##### Amplification

Amplification was required to copy the cDNA off the beads, and was conducted using the PCR handle on round_3_barcoding oligos and the newly added handle from second strand synthesis. After the final water wash following second strand synthesis, beads were resuspended in 220 μL PCR mix per library. Each mix was distributed across 4-5 wells of a PCR 96-well plate and run with the following thermocycler program: 95°C for 3 min, 3 cycles of [98°C for 20 sec, 65°C for 45 sec, 72°C for 3 min]. The plate was then removed from the thermocycler, mixed by pipetting to resuspend settled beads, and returned to the thermocycler to repeat the same program with 2 cycles. After a total of 5 cycles, the plate was removed, briefly mixed by pipetting, and placed onto a 96-well plate magnetic stand where the beads were pulled out of solution and discarded. The remaining solution was moved into new wells for three consecutive SPRIselect bead cleanups: 0.90X with elution in 100 μL water, 0.80X with elution in 50 μL water, and 0.80X with final elution in 21 μL water. Separate wells for each library were combined back into a single well during elution in the first cleanup. Throughout cell lysis, purification, and second strand synthesis, there was a greater amount of free/unbound round_3_barcoding oligos in solution than RNA constructs. These free/unbound round_3_barcoding oligos were biotinylated, which resulted in their capture on the streptavidin magnetic beads alongside the desired RNA constructs. If left in solution for too many amplification cycles after second-strand synthesis, erroneous PCR combination events leading to long polymer products were observed (Figure S1C). Since the problematic round_3_barcoding oligos were shorter than RNA constructs (which become double-stranded cDNA after second strand synthesis), they could be removed based on their size by SPRIselect bead cleanups. Three consecutive cleanups were performed as described to eliminate the large excess of small products, and the elution volumes were decreased sequentially to prevent losses from using a small volume to elute from a large amount of beads. After the initial 5 amplification cycles and cleanup, the final 21 μL of eluted product was combined with 2 μl each of PCR_P1 and PCR_P2 primers (10 μM each) and 25 μL 2X KAPA HiFi HotStart ReadyMix and placed in the thermocycler for additional amplification with the following program: 95°C for 3 min, 15-18 cycles of [98°C for 20 sec, 67°C for 20 sec, 72°C for 3 min], 72°C for 5 min. Amplification was performed until reaching a concentration of 6-10 ng/μL measured by Qubit, which was generally achieved in 20-23 total cycles (including the initial 5 cycles). After each set of cycles, a single SPRIselect bead cleanup was performed at 0.80X with elution in 21 μL water. After reaching this target concentration, fragment analysis was used to confirm a quality amplification product with fragment sizes ranging from 300-3,000 bp (Figure S1B).

##### Note on library concentrations

A target library concentration of 6-10 ng/μL was sufficient for downstream processing with reserve in case there was a need to return to the previous step. Moving through each step following amplification, we ensured at least 25% of the library volume was carried forward to ensure representation of full library complexity.

#### rRNA depletion: Dual-strand subtractive hybridization

##### Forward and reverse oligo master mixes

Sets of biotinylated oligos designed against 23S, 16S, and 5S fragments for rRNA depletion from double-stranded cDNA (rRNA_dep_Fwd and rRNA_dep_Rev) were each suspended at 100 μM in water from the solid desalted form. An equal volume (5 μL) of all 21 oligos from forward or reverse sets were pooled into a single master mix for either forward (MM-Fwd) or reverse (MM-Rev) strand depletion, being careful not to combine forward and reverse oligos. In the pooled mixes with equal volume of 21 forward or reverse oligos, the concentration of each oligo was 4.76 μM.

##### Forward and reverse depletion mixes

Working mixes for depletion of forward and reverse rRNA fragments (dep-M-Fwd and dep-M-Rev) were comprised of SSC buffer (20X), EDTA (100 mM), T.1E, and MM-Fwd or MM-Rev. The specific amounts of each component was calculated using the Excel calculator tool developed by Culviner et al. based on total RNA input, desired oligo:RNA and bead:oligo ratios, and total reaction volume.^30^ In our implementation of dual-strand subtractive hybridization, total RNA input was variable based on the amplified library concentration following second strand synthesis (generally in the range of 200-250 ng), oligo:RNA ratio was set at 5, bead:oligo ratio was set at 18, and the reaction volume was set at 70 μL. In the calculator tool, the number of oligos should be set at 21 and the bead capacity should be set to 1.60×10^−12^ mol/μL if using NEB Hydrophilic Streptavidin Magnetic Beads.

<u>Example</u> – An amplified library after second strand synthesis is 29 µL at a concentration of 8.28 ng/μL. If nearly the full volume is used, then the total RNA input can be set at 230 ng (requiring 27.8 μL of the library), oligo:RNA=5, bead:oligo=18, and reaction volume at 70 μL. The forward depletion mix (dep-M-Fwd) is made according to the calculator tool^30^ as 43.75 μL 20X SSC, 8.75 μL 100 mM EDTA, 3.67 μL MM-Fwd, and 124.4 μL T.1E. Note that the total volume of dep-M-Fwd is about 25X larger than what is required; however, the large volume is used such that the volume of MM-Fwd is accurately measurable. To calculate the requirements for the reverse depletion mix (dep-M-Rev), the theoretical concentration in 25 μL is input as the concentration (230 ng / 25 μL = 9.2 ng/μL) with all other settings held uniform. This is because depletion of the reverse strand is initiated with 25 μL volume. The dep-M-Rev mix is made as 43.75 μL 20X SSC, 8.75 μL 100 mM EDTA, 3.67 μL MM-Rev, and 193.83 μL T.1E. Again the total volume of dep-M-Rev is about 25X larger than what is required.

##### Prepare streptavidin beads

The volume of streptavidin beads was also determined by the calculator tool. Since our method depleted from both the forward and reverse strands, the amount of beads was doubled.

<u>Example (continued from above)</u> – 331 μL of beads are used. The beads are washed three times with ∼500 μL of 5X SSC, then resuspended in 70 μL 5X SSC and kept at room temperature until used.

##### Forward strand depletion and cleanup

The library was first denatured in a thermocycler by heating to 95°C for 2 min and then cooling to 20°C. A 96-well PCR plate was used for all depletion steps. Then, the dep-M-Fwd mix was added as directed in the calculator tool to achieve a final volume of 35 μL. Annealing of the depletion oligos to cDNA template was carried out in a thermocycler with the following program: 95°C for 3 min, 98°C for 20 sec, 70°C for 5 min, decrease to 20°C at a rate of −0.1°C/sec.

Example (continued from above) – 27.8 μL of library is denatured, then 7.2 μL of dep-M-Fwd is added and run on the annealing program.

Following annealing, the rRNA constructs were depleted by adding 35 μL washed streptavidin beads and mixing by pipetting at least 25X. The bead mixture was incubated at room temperature with shaking (∼500-700 rpm) and intermittent mixing with a pipette for 10 min. After a final pipette mixing step, bead mixtures were incubated in a thermocycler at 50°C for 5 min (lid temperature reduced to 50°C). Mixtures were then immediately set on a magnetic plate and the supernatants transferred to new wells after the beads had adequately settled (beads were discarded). SPRIselect bead cleanup was then performed on the supernatants at 0.90X with elution in 25 μL water.

##### Reverse strand depletion and cleanup

The 25 μL volume from forward strand depletion and cleanup was heated to 95°C for 2 min and then cooled to 20°C. Then, 10 μL dep-M-Rev mix was added, and annealing of the depletion oligos to cDNA template was performed in a thermocycler with the same annealing program described for forward strand depletion. As for the forward strand, 35 μL of washed streptavidin beads were added and incubated for 10 min at room temperature with mixing before heating at 50°C for 5 min and removing the beads via magnet. A 0.90X SPRIselect bead cleanup was then performed with elution in 21 μL water.

##### Amplification

The final eluted 21 μL of rRNA-depleted product was combined with 2 μl each of PCR_P1 and PCR_P2 primers (10 μM each) and 25 μL 2X KAPA HiFi HotStart ReadyMix and amplified with the following thermocycler program: 95°C for 3 min, 4-5 cycles of [98°C for 20 sec, 67°C for 20 sec, 72°C for 3 min], 72°C for 5 min. Amplification was performed until reaching a concentration of 10-20 ng/μL measured by Qubit, which was generally achieved in 4-5 total cycles. A 0.90X SPRIselect bead cleanup was then performed with elution in 21 μL water. Fragment analysis was used to confirm a quality amplification product with fragment sizes ranging from 300-3,000 bp (Figure S1B). In comparison to the size distributions following second strand synthesis, size distributions after rRNA depletion were shifted towards the 700-3,000 bp range. As an additional quality check, qPCR was performed side-by-side on samples before and after depletion. Analysis of 23S and 16S rRNA abundance in relation to a control gene (*gyrB*) showed substantial rRNA depletion with minimal alterations to the control gene (Figure S1D). The qPCR was performed with Luna Universal qPCR Master Mix from NEB and primer sets for *gyrB* (gyrB_Fwd and gyrB_Rev), 23S rRNA (23S_Fwd and 23S_Rev), and 16S rRNA (16S_Fwd and 16S_Rev).

#### Library prep and sequencing

##### Fragmentation and cleanup

Reagents used in fragmentation were from the sparQ DNA fragment and library prep kit (Quantabio). A mix of 5X DNA Frag & Polishing Enzyme Mix (10 μL per library) and 10X DNA Frag & Polishing Buffer (5 μL per library) was first prepared and kept on ice. Next, 15 μL of this mix was added to 150-250 ng rRNA-depleted sample library in 35 μL water, for a total volume of 50 μL which was also held on ice. The 50 μL fragmentation mix was quickly transferred to a pre-chilled thermocycler and run on the following program: 4°C for 1 min, 32°C for 8 min (fragmentation step), 65°C for 30 min, and 4°C hold. The samples were immediately removed and stored on ice before quickly initiating a double-side 0.825-0.45X SPRIselect bead cleanup. Final product was eluted in 30 μL water, then brought up to 50.5 μL with water.

##### Ligation

Reagents used in ligation were also from the sparQ DNA fragment and library prep kit (Quantabio), with additional custom oligos. A mix of 5X DNA Rapid Ligation Buffer (20 μL per library), DNA Ligase (10 μL per library), and water (17.5 μL per library) was prepared and kept on ice. The adapter duplex consisting of adapter_duplex_Top and adapter_duplex_Bott was pre-annealed by heating an adapter mix (100 μM each in 50 μM NaCl) to 95°C and cooling to 20°C at a rate of −0.1°C/sec. Then, 2 μL of the pre-annealed adapter duplex mix was added to each 50.5 μL library (after double-sided bead cleanup) on ice. Finally, 47.5 μL of the ligation mix was added to each 52.5 μL library with adapters, and the 100 μL solution was added to a thermocycler for incubation at 20°C for 15 min (with unheated lid). The ligation reactions were immediately removed from the thermocycler and SPRIselect cleaned two consecutive times at 0.81X, first eluting in 50 μL water and then 21 μL water on the final cleanup.

##### Amplification

The 21 μL cleaned ligation product was combined with 2 μL each of i5 primer (seq_i501/2) and i7 primer (seq_i701/2) (10 μM each) and 25 μL 2X KAPA HiFi HotStart ReadyMix and amplified with the following thermocycler program: 95°C for 3 min, 4-6 cycles of [98°C for 20 sec, 67°C for 20 sec, 72°C for 3 min], 72°C for 5 min. The number of cycles depended on the input amount, where 250 ng input required only 4 cycles, whereas 150 ng input required 6 cycles. Next, two consecutive double-sided SPRIselect bead cleanups were performed, first at 0.75-0.45X and then at 0.775-0.425X. Two cleanups were necessary to adequately remove fragments that were too small or large. Fragment analysis was performed for final library quality control (Figure S1B).

##### Library quantification

NEBNext Library Quant Kit for Illumina was used for qPCR-based library quantification (Figure S1E). With the described library prep protocol, final libraries were in the range of 1-2 nM. We found it beneficial to keep library concentrations low to maximize the amount of each library loaded on the sequencer, which ensured capturing full diversity within the library.

##### Sequencing

The datasets presented were sequenced on an Illumina NextSeq 550 series sequencer with v2.5 150-cycle mid-output kit. Read 1 (insert/transcript) was allocated 63 cycles, and read 2 (barcodes) was allocated 89 cycles, 3 extra bases than the full length barcodes that are 86 bp.

#### Count matrix generation

##### Remove adapters and quality filter

Cutadapt^169^ was used to identify and remove sequencing adapters from the ends of read 1 and read 2 (AGATCGGAAGAGCACACGTCTGAACTCC and ACTGTCTCTTATACACATCT) and filter out reads with quality scores <10.

##### Extract UMIs

UMI sequences were extracted from read 2 with UMI-tools^170^ using the *extract* command.

##### Demultiplexing

Cutadapt was used to demultiplex reads by the 5’-end into unique combinations of three barcodes beginning with barcode 3, then barcode 2, and barcode 1. A python script was used to loop Cutadapt through all files, and each unique barcode combination was output into a separate file. Barcode 3 was demultiplexed as the 8-nucleotide barcode and linker sequence between barcodes 2 and 3 (totaling 38 nucleotides), and processed as a non-internal 5’ adapter with minimum overlap of 35 and error tolerance set at 0.15. Barcode 2 was demultiplexed as the 8-nucleotide barcode and linker sequence between barcodes 1 and 2 (totaling 30 nucleotides), and processed as a non-internal 5’ adapter with minimum overlap of 25 and error tolerance set at 0.2. Barcode 1 was demultiplexed as the 8-nucleotide barcode and processed as a non-internal 5’ adapter with minimum overlap of 6. After demultiplexing, cells were separated by sample based on barcode 1.

##### Alignment

After demultiplexing, only read 1 sequences were maintained for processing. Reads were mapped to the annotated *S. aureus* USA300 FPR3757 genome (NCBI)^171^ using STAR^172^. As previously reported, we ignored splicing detection and retained multimapping reads, as bacterial genomes are known to contain overlapping coding sequences.^20^ Additionally, many reads had either full or partial fragments of the second strand synthesis oligo (S^3^_randomer, 34 nucleotides), which reduced the portion of the read available for mapping. Therefore, the options *-outFilterMatchNminOverLread* and *-outFilterScoreMinOverLread* were reduced to 0.2 to increase tolerance for smaller fractions of the aligning read. This adjustment was necessary to prevent loss of transcriptional information and data bias; however, it also led to inherent background noise as extraneous multimapping reads were inevitably identified.

##### Counting

The featureCounts^173^ package of Subread was used to annotate and enumerate transcript features after alignment. Multimapping reads were assigned a fractional count, which was necessary for accurately quantifying rRNA reads that aligned to multiple regions in the genome, as well as to attenuate background noise from spurious alignments.

##### Collapse UMIs

The *dedup* command within UMI-tools was used to deduplicate reads based on UMI and mapping coordinates.

##### Count matrix

A matrix of cells-by-genes was filled with gene counts per cell, for input into Scanpy.

#### Data processing and clustering

##### Preprocessing, filtering, and normalization

Data analysis was performed in Scanpy.^174^ All rRNA genes were removed, and cells were filtered based on total mRNA and tRNA (non-rRNA) expression levels. For the biofilm vs. planktonic comparison experiment, biofilm cells having ≥7 and planktonic cells having ≥28 non-rRNA reads per cell were maintained for analysis, totaling 3,680 and 4,231 cells respectively (Figure S3A). Resulting rRNA, mRNA, and tRNA metrics are shown in Figure S3B-C. For the biofilm-leukocyte co-culture experiment, cells from all conditions were kept for analysis if they contained ≥15 non-rRNA reads per cell, totaling 4,655, 4,544, 4,780, and 6,125 cells for the biofilm control, biofilm + MΦs, biofilm + G-MDSCs, and biofilm + PMNs respectively (Figure S5A). Resulting rRNA, mRNA, and tRNA metrics are shown in Figure S5B-C. All cells were uniformly normalized to 10^4^ total counts and log+1 transformed.

##### Clustering

For the biofilm and planktonic comparison experiment, cells from each growth state were combined and principal components calculated from highly variable genes (*min_mean*=0.00625 and *min_disp*=0.25). Nearest neighbors were determined and integrated with batch balanced k nearest neighbors (BBKNN) alignment (*neighbors_within_batch*=9 and *n_pcs*=4).^60^ Neighbors were UMAP embedded (*min_dist*=0.24 and *spread*=0.21) and clustered with the Leiden algorithm (*resolution*=0.205). For the biofilm-leukocyte co-culture experiment, cells from each condition were first independently analyzed with principal components calculated from highly variable genes (*min_mean*=0.00625 and *min_disp*=0.25). Nearest neighbors were determined (*n_neighbors*=12 and *n_pcs*=7), UMAP embedded (*min_dist*=0.5 and *spread*=1), and clustered with the Leiden algorithm (*resolution*=0.15). For comparison of biofilm responses to MΦs, G-MDSCs, and PMNs, biofilm cells from each condition were projected onto the UMAP of the biofilm control using the *ingest* function of Scanpy. The noted clustering settings were chosen to optimize resolved sets of marker genes, which were used as a quality control readout (Figure S3D and Figure S6).

##### Differential expression

The MAST algorithm^61^ was integrated with the Scanpy workflow through the rpy2, anndata2ri, and sc_toolbox packages and IPython interface in Jupyter notebook.^175^ MAST was used for all differential expression analyses. A known caveat of MAST is that log_2_ fold-change values can be small; therefore, marginal differences cannot be disregarded as insignificant.

#### iModulon and transcriptional regulatory network analyses

Gene sets were downloaded from the *S. aureus* Precise165 dataset on the iModulonDB database^36^ as determined in Poudel et al.,^34^ and USA300_TCH1516 annotations were converted to USA300_FPR3757 orthologues using AureoWiki databases.^176^ On a per cell basis, raw gene expression was summed for all genes within each iModulon, forming an expression score for each iModulon. Then, raw iModulon scores were summed for all iModulons comprising a transcriptional regulatory network category, forming a regulatory category score. The iModulon and transcriptional regulatory network scores were uniformly normalized to 10^4^ total counts and log+1 transformed separately from the individual genes.

##### Trajectory analysis

The Palantir package was used for trajectory analysis.^37^ Diffusion maps (*n_components*=5) were first determined, and MAGIC imputation^97^ (from within the Palantir package) was used for resolving expression trends. Trajectory analysis (*num_waypoints*=500) was then performed with an indicated starting cell and terminal states, and importantly altering the cell of origin did not affect trajectory outcome. Detailed initiating and terminal state selections, and entropy quantifications are shown in Figure S4. Pearson correlations were calculated for the imputed expression of each gene over the Palantir-calculated pseudotime variable. Top lists of positively and negatively correlated genes with pseudotime are shown in Tables S1-S2.

### STATISTICAL ANALYSES

The statistical tests for all BaSSSh-seq experiments were performed in python 3 (3.10.12) within Jupyter notebook with the respective analysis or GraphPad Prism (10.0.2). Statistical details can be found in figures and legends where appropriate.

