## Supplemental Information for "Bacterial single-cell RNA sequencing captures biofilm transcriptional heterogeneity and differential responses to immune pressure"

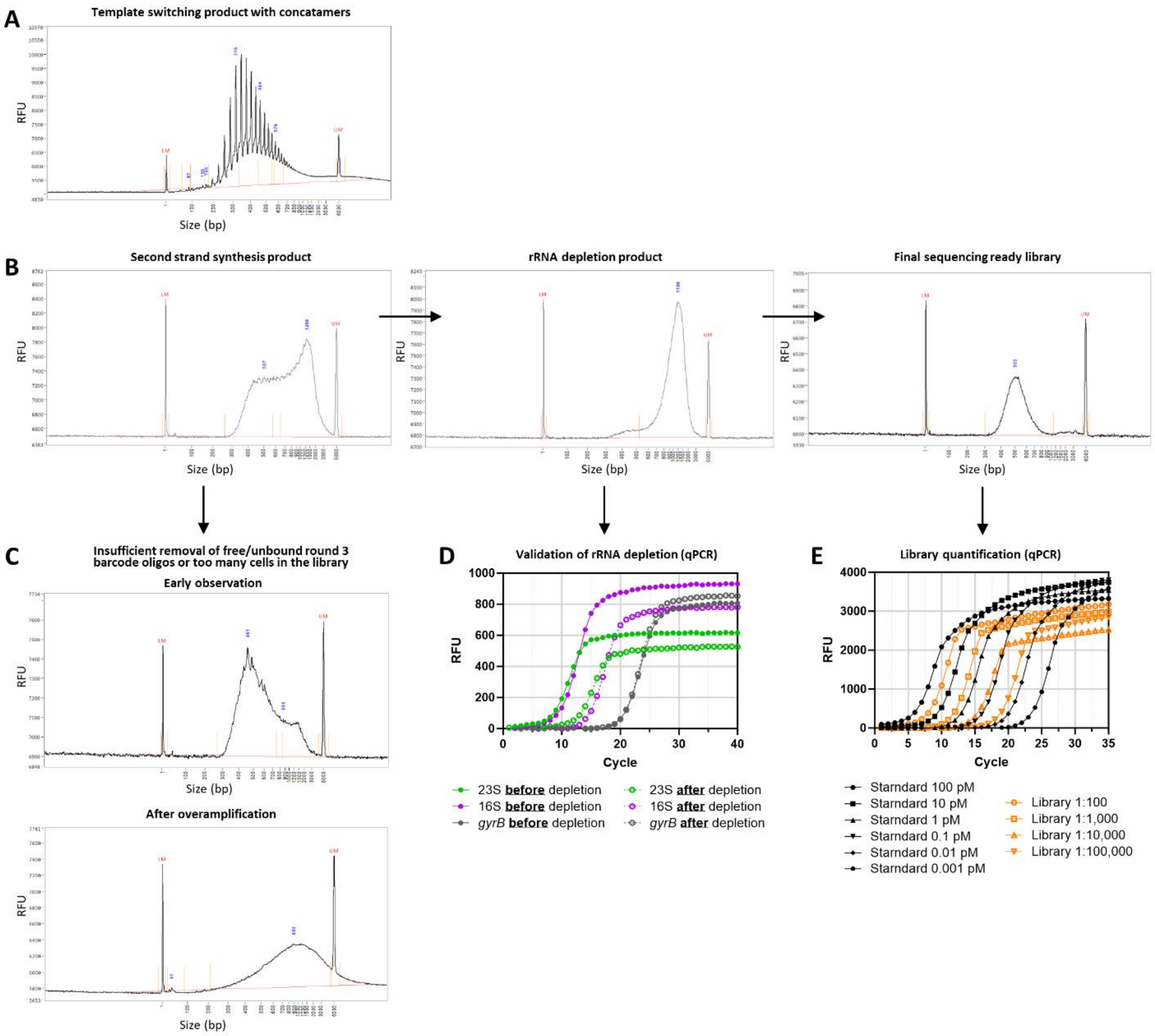


**Figure S1. BaSSSh-seq quality control steps**

(A) Representative size distribution from fragment analysis on a template switching product. Concatamers are seen as equally spaced spikes across the size distribution, with spikes corresponding to the length of the template switching oligo used. The raised baseline extending past the upper 6,000 bp marker is also reflective of concatamer products.

(B) Representative size distributions from fragment analysis during each of the quality control checks in the BaSSSh-seq process. Check points occur after second strand synthesis (left), rRNA depletion (middle), and library prep (right).

(C) Examples of size distributions from fragment analysis after second strand synthesis if there was insufficient removal of free/unbound round 3 barcode oligos or too many cells in the library. If amplification is stopped early, the size distribution is biased towards smaller fragments, and if amplification is prolonged the size distribution becomes larger and raises the baseline past the upper 6,000 bp marker.

(D) Representative validation of rRNA depletion by qPCR. Sample aliquots from before and after rRNA depletion were analyzed for 23S and 16S rRNA abundance in relation to *gyrB* as a control. Curves were used to calculate $2^{-\Delta\Delta Cq}$ values for each sample (0.038 and 0.026 for 23S and 16S, respectively in the representative example).

(E) Representative library quantification by qPCR. Libraries were diluted and analyzed in comparison to a standard curve.

For (A-C), all size distributions were measured on a 5200 Fragment Analyzer (Agilent). Lower markers (LM) are indicated at 1 bp, and upper markers (UM) are indicated at 6,000 bp.

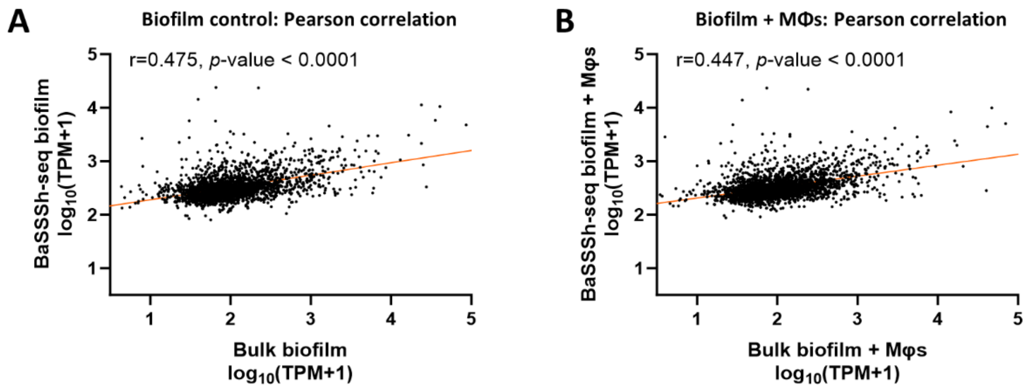


**Figure S2. Correlation of BaSSSh-seq with bulk RNA-seq**

Transcriptomic profiles from single-cell RNA-seq with BaSSSh-seq significantly correlate with those captured in a prior report (doi: 10.1128/iai.00428-22) using bulk RNA-seq for biofilm control (no immune cells) (A) and biofilm +MΦs co-culture (B).

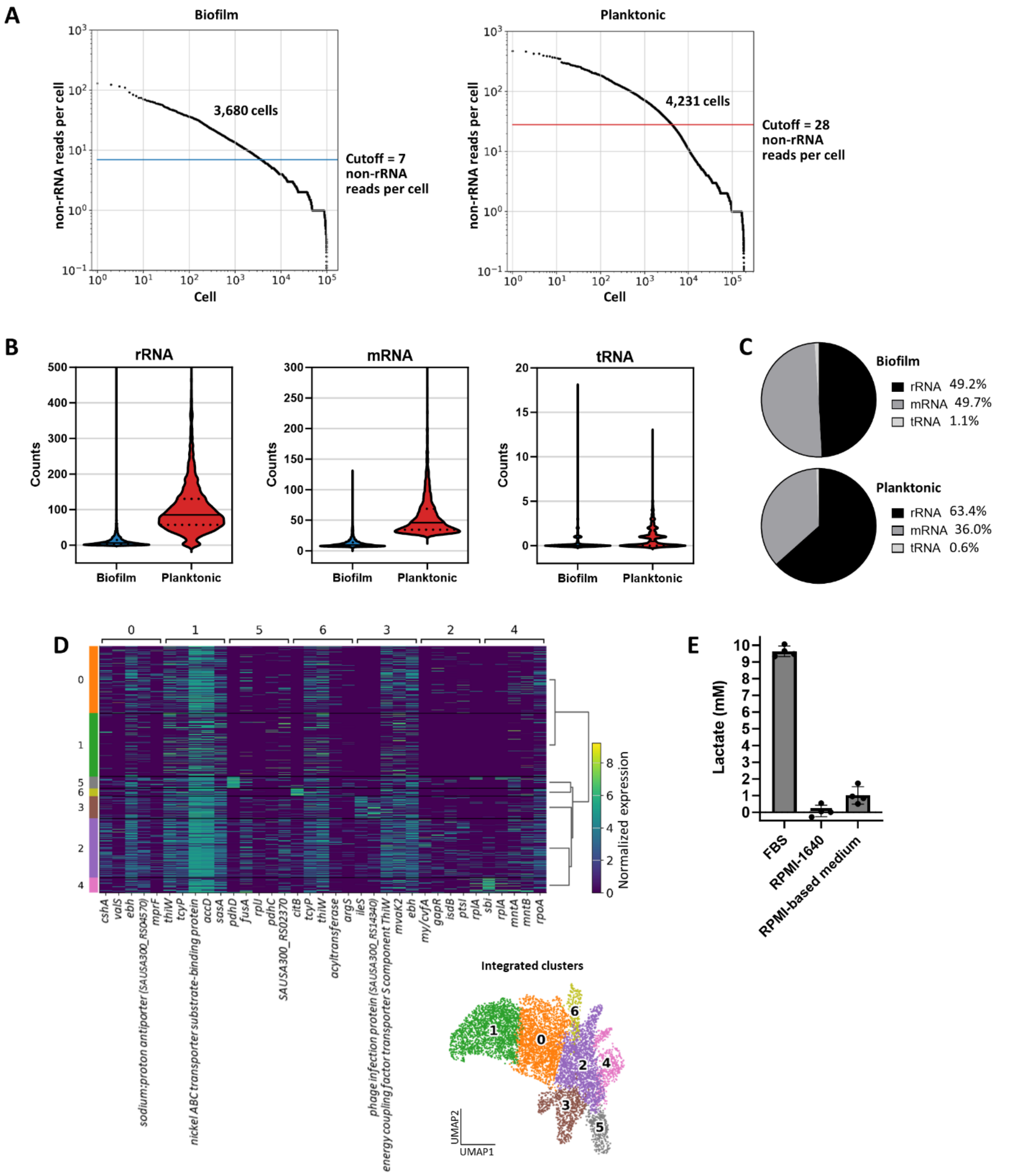


**Figure S3. Filtering, metrics, and clustering when comparing biofilm and planktonic growth with BaSSSh-seq**

(A) Filtering sequenced cells based on the number of non-rRNA (mRNA and tRNA) reads per cell. Cells are sorted in terms of decreasing reads per cell. Numbers of cells carried through for analysis are noted on the plots.

(B) Counts of rRNA, mRNA, and tRNA for filtered cells from biofilm and planktonic samples. Solid lines indicate the median, and dotted lines reflect upper and lower quartiles.
(C) Percentages of rRNA, mRNA, and tRNA for filtered cells from biofilm and planktonic samples.
(D) Marker gene heatmap arranged by dendrogram relation for the integrated biofilm and planktonic samples, with UMAP on the bottom right (UMAP originally defined in Figure 1C).
(E) Lactate levels in fetal bovine serum (FBS), RPMI-1640, and the RPMI-based medium used for *S. aureus* culture containing 10% FBS.

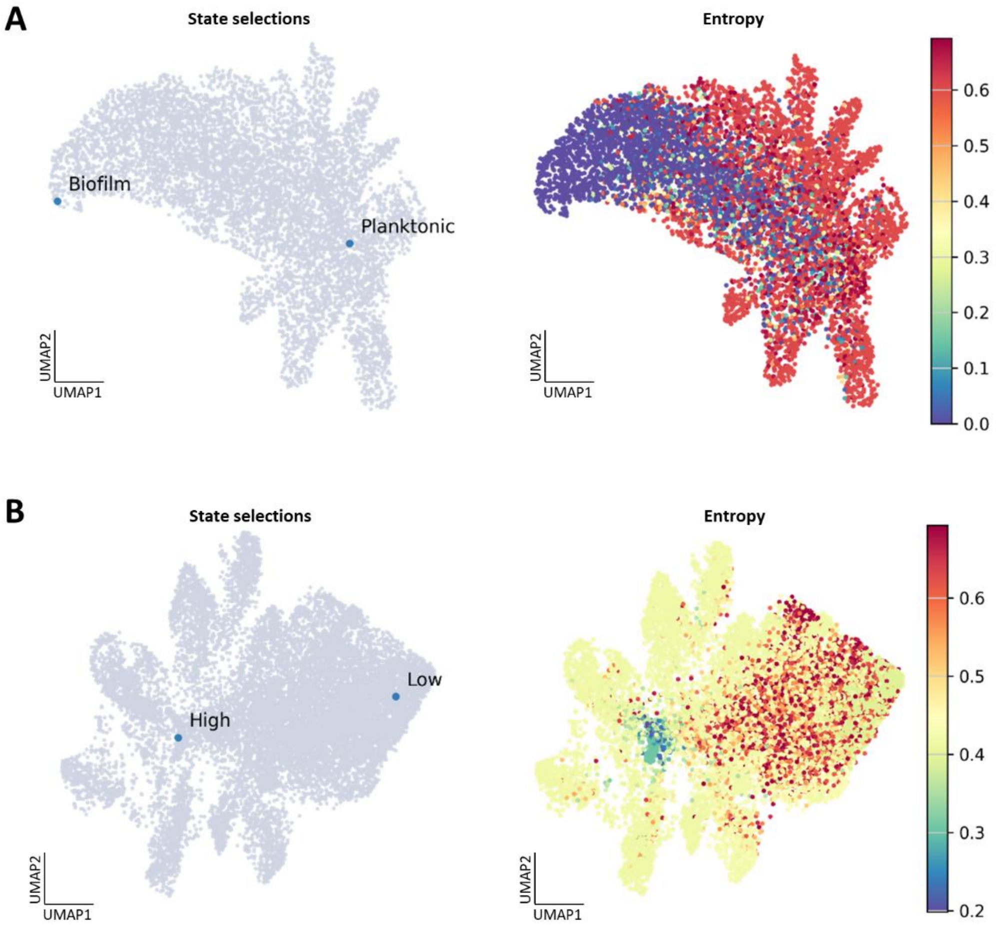


**Figure S4. Trajectory analysis details**

(A)Trajectory details for biofilm and planktonic growth comparison experiment. (Left) Cell selections denoting the *terminal_states* parameters in the Palantir trajectory algorithm, for both 'Biofilm' and 'Planktonic'. (Right) Entropy plot for the integrated biofilm and planktonic samples.

(B) Trajectory details for biofilm-leukocyte co-culture experiment. (Left) Cell selections denoting the *terminal_states* parameters in the Palantir trajectory algorithm, selected to represent 'High' activity and 'Low' activity. (Right) Entropy plot for the biofilm-leukocyte co-culture samples. The trajectories for both experiments (Figure 3E and Figure 6C) were robust to changes in the *terminal_states* parameter selections.

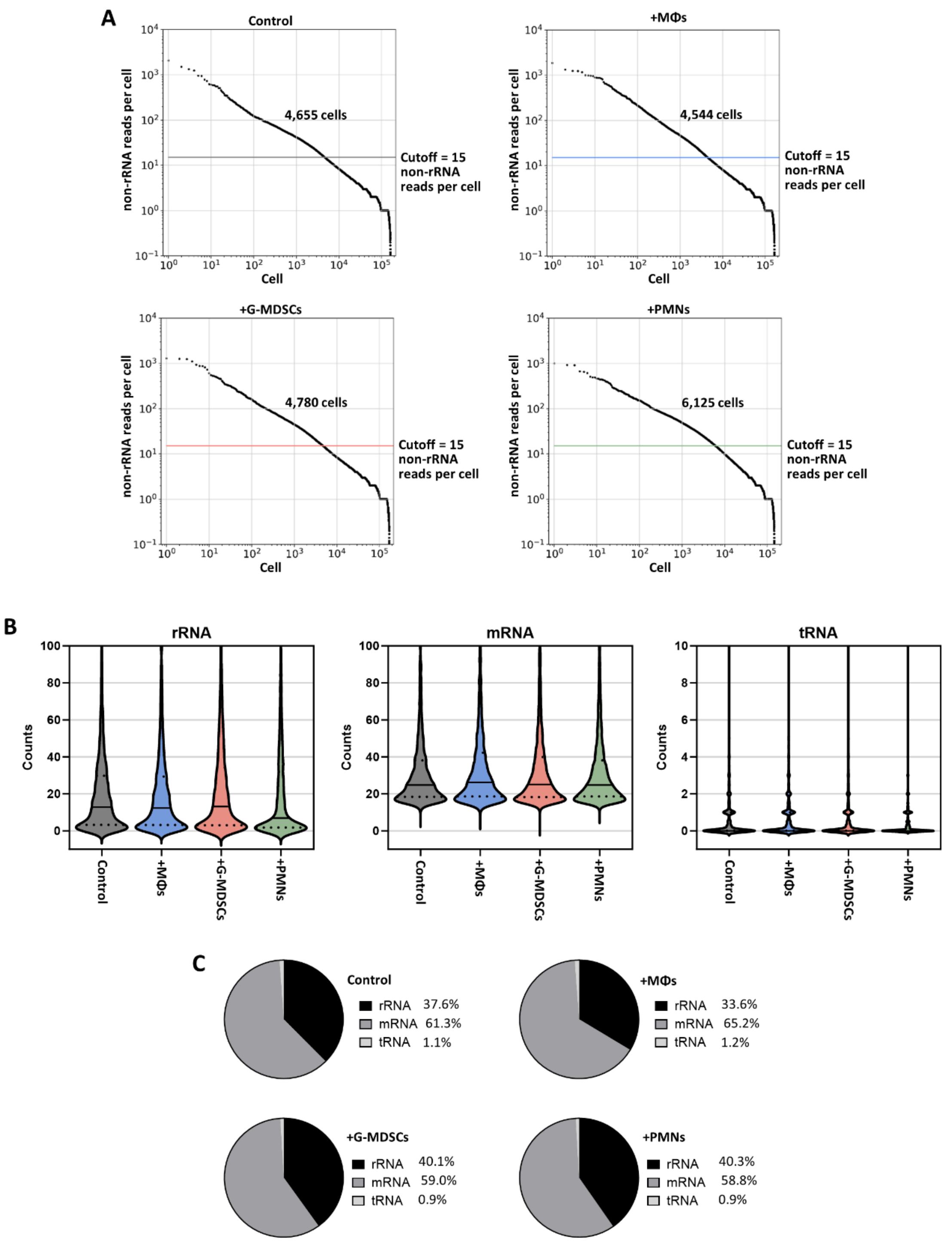


**Figure S5. Filtering and metrics for biofilm-leukocyte co-culture experiments with BaSSSh-seq**

(A) Filtering sequenced cells based on the number of non-rRNA (mRNA and tRNA) reads per cell. Cells are sorted in terms of decreasing reads per cell. Numbers of cells carried through for analysis are noted on the plots.

(B) Counts of rRNA, mRNA, and tRNA for filtered cells from co-culture samples. Solid lines indicate the median, and dotted lines reflect upper and lower quartiles.
(C) Percentages of rRNA, mRNA, and tRNA for filtered cells from co-culture samples.

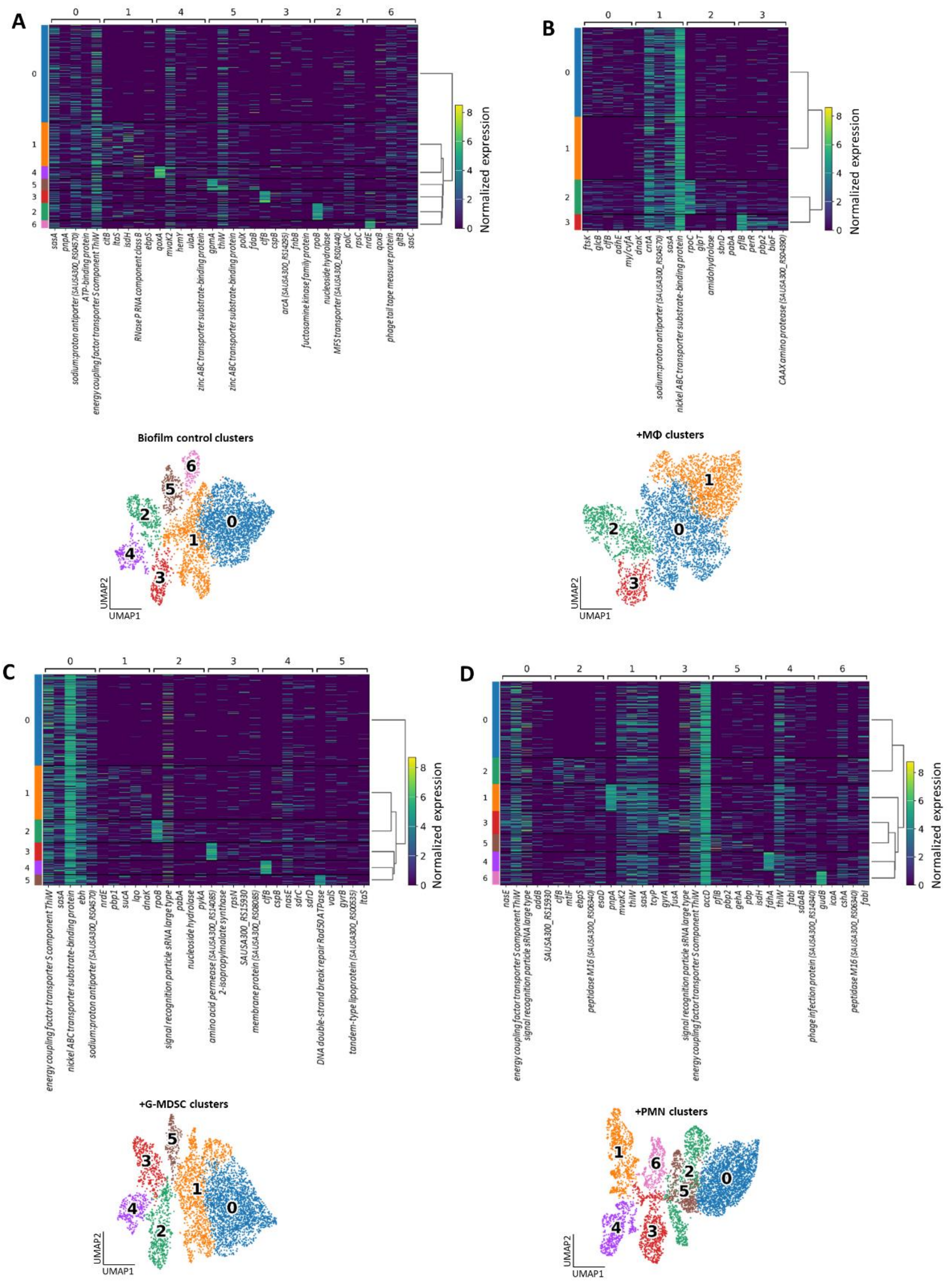


**Figure S6. Clustering for biofilm-leukocyte co-culture experiments with BaSSSh-seq**

Marker gene heatmaps arranged by dendrogram relation for the biofilm control (A), and independently clustered biofilm + MΦs (B), biofilm + G-MDSCs (C), and biofilm + PMNs (D), with UMAPs below (UMAPs also shown in Figure 4B-C). For comparative analysis, all co-cultured biofilm cells were projected onto the control UMAP in Figure 6.

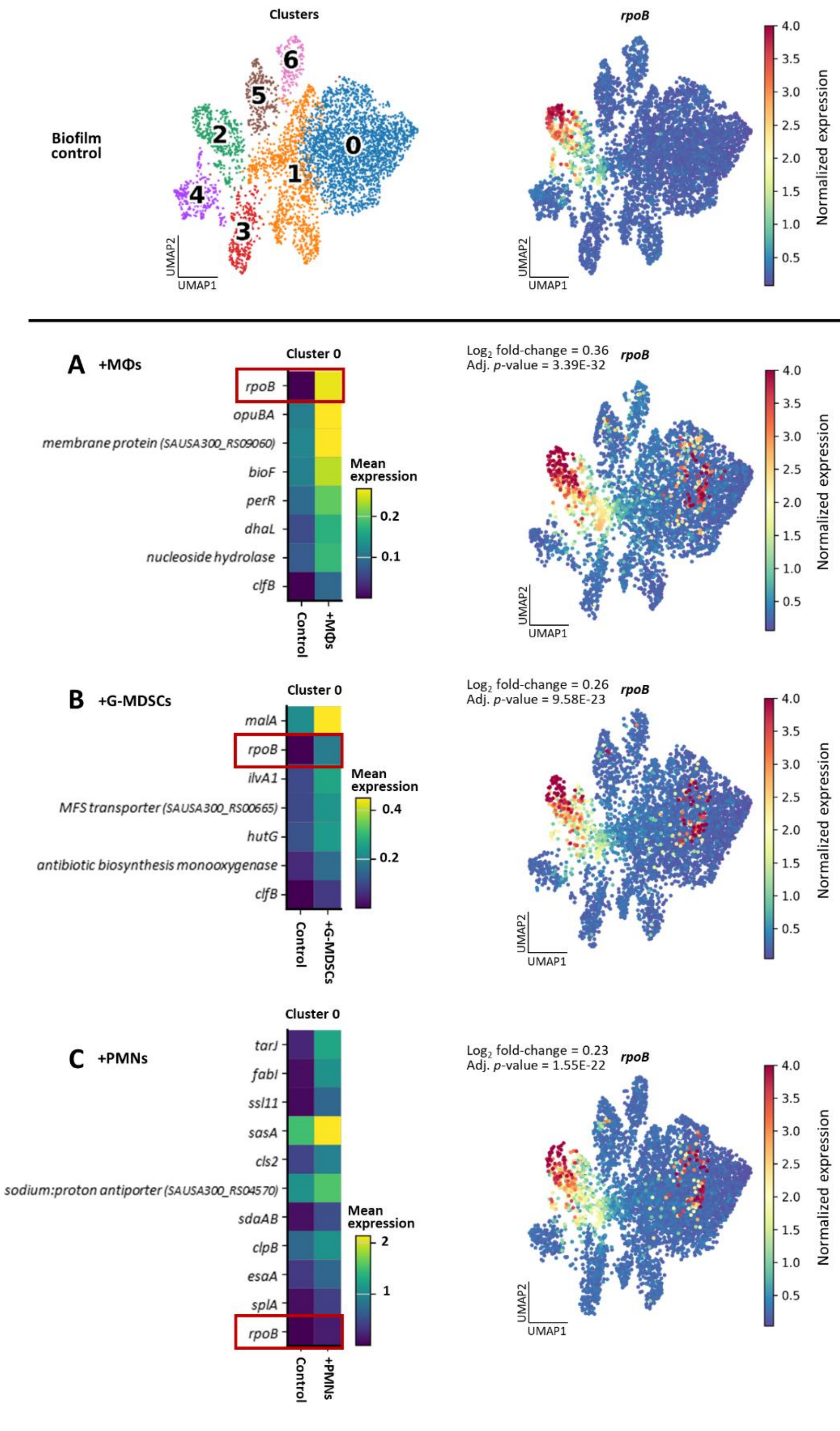


**Figure S7. Transcriptional evidence of a potential biofilm persister cell population**

Differential expression was performed using the MAST algorithm for cluster 0 on each of the biofilm-leukocyte co-cultures compared to the biofilm control. Each co-culture condition displayed a set of genes significantly upregulated within cluster 0, a small subset of which are displayed in the matrix plots for +MΦs (A), +G-MDSCs (B), or +PMNs (C). RNA polymerase (*rpoB*) was consistently upregulated across each condition, which is further illustrated in UMAP overlays of *rpoB* expression displayed to the right of each matrix plot, with the biofilm control shown at the top along with cluster identities for reference.

### Table S1. Genes positively correlated with the trajectory in Figure 3E

| Gene symbol/descriptor | Locus tag | TIGRFAM main role | TIGRFAM sub role | Pearson correlation | *p*-value |
|---|---|---|---|---|---|
| - | SAUSA300_RS15665 | - | - | 0.447 | <1.0E-323 |
| - | SAUSA300_RS07700 | - | - | 0.432 | <1.0E-323 |
| *psmβ1* | SAUSA300_RS05790 | Virulence | Virulence | 0.385 | 3.5E-278 |
| - | SAUSA300_RS11765 | - | - | 0.381 | 2.9E-271 |
| - | SAUSA300_RS07610 | - | - | 0.360 | 4.1E-241 |
| - | SAUSA300_RS07460 | - | - | 0.350 | 2.4E-226 |
| - | SAUSA300_RS15565 | - | - | 0.333 | 4.0E-204 |
| *cwrA* | SAUSA300_RS13850 | - | - | 0.332 | 2.0E-202 |
| - | SAUSA300_RS05335 | - | - | 0.331 | 4.9E-202 |
| - | SAUSA300_RS07785 | - | - | 0.328 | 9.3E-198 |
| - | SAUSA300_RS06535 | - | - | 0.315 | 2.0E-181 |
| - | SAUSA300_RS07605 | - | - | 0.311 | 5.1E-177 |
| - | SAUSA300_RS01575 | - | - | 0.302 | 4.0E-166 |
| *membrane protein* | SAUSA300_RS14650 | Cell envelope | Other | 0.284 | 4.2E-147 |
| *esaG* | SAUSA300_RS01545 | Protein fate | Protein and peptide secretion and trafficking | 0.273 | 1.5E-135 |
| - | SAUSA300_RS06540 | - | - | 0.273 | 2.0E-135 |
| - | SAUSA300_RS11950 | - | - | 0.235 | 9.6E-100 |
| - | SAUSA300_RS15730 | - | - | 0.223 | 5.4E-90 |
| *aroB* | SAUSA300_RS07390 | Amino acid biosynthesis | Aromatic amino acid family | 0.219 | 1.4E-86 |
| *estA* | SAUSA300_RS14265 | Cellular processes | Detoxification | 0.203 | 4.2E-74 |
| - | SAUSA300_RS11910 | - | - | 0.199 | 8.5E-72 |
| - | SAUSA300_RS07790 | - | - | 0.197 | 1.0E-69 |
| - | SAUSA300_RS09870 | - | - | 0.194 | 4.9E-68 |
| - | SAUSA300_RS13255 | - | - | 0.193 | 2.4E-67 |
| *TIGR01741 family protein* | SAUSA300_RS01600 | - | - | 0.181 | 3.2E-59 |
| *trnaG* | SAUSA300_RS09915 | - | - | 0.181 | 4.4E-59 |
| *HNH endonuclease* | SAUSA300_RS10660 | - | - | 0.178 | 4.7E-57 |
| *protein VraX* | SAUSA300_RS03005 | - | - | 0.173 | 2.9E-54 |
| - | SAUSA300_RS05675 | - | - | 0.170 | 3.0E-52 |
| - | SAUSA300_RS10405 | - | - | 0.169 | 9.2E-52 |
| - | SAUSA300_RS13250 | - | - | 0.162 | 2.1E-47 |
| - | SAUSA300_RS15375 | - | - | 0.161 | 4.5E-47 |
| *peroxiredoxin* | SAUSA300_RS10070 | Cellular processes | Detoxification; Adaptations to atypical conditions | 0.154 | 2.4E-43 |
| - | SAUSA300_RS11530 | - | - | 0.146 | 7.7E-39 |
| *TIGR01741 family protein* | SAUSA300_RS01605 | - | - | 0.146 | 9.7E-39 |
| *ftnA* | SAUSA300_RS10250 | Transport and binding proteins | Cations and iron carrying compounds | 0.145 | 3.1E-38 |
| - | SAUSA300_RS15905 | - | - | 0.140 | 4.4E-36 |
| - | SAUSA300_RS11700 | - | - | 0.139 | 2.0E-35 |
| - | SAUSA300_RS10285 | - | - | 0.138 | 3.6E-35 |
| *rimM* | SAUSA300_RS06125 | Transcription | RNA processing | 0.137 | 3.1E-34 |
| - | SAUSA300_RS05495 | - | - | 0.132 | 2.8E-32 |
| - | SAUSA300_RS07760 | - | - | 0.132 | 6.2E-32 |
| *rpoY* | SAUSA300_RS05330 | - | - | 0.129 | 8.1E-31 |
| - | SAUSA300_RS06565 | - | - | 0.126 | 1.8E-29 |
| *cysteine hydrolase* | SAUSA300_RS14350 | - | - | 0.122 | 1.4E-27 |

### Table S2. Genes negatively correlated with the trajectory in Figure 3E

| Gene symbol/descriptor | Locus tag | TIGRFAM main role | TIGRFAM sub role | Pearson correlation | *p*-value |
|---|---|---|---|---|---|
| *cntA* | SAUSA300_RS13350 | Transport and binding proteins | Cations and iron carrying compounds | -0.819 | <1.0E-323 |
| *fusA* | SAUSA300_RS02845 | Protein synthesis | Translation factors | -0.816 | <1.0E-323 |
| *energy coupling factor transporter S component ThiW* | SAUSA300_RS06415 | Biosynthesis of cofactors, prosthetic groups, and carriers; Transport and binding proteins | Thiamine; Other | -0.812 | <1.0E-323 |
| *ATP-binding protein* | SAUSA300_RS09575 | DNA metabolism | DNA replication, recombination, and repair | -0.798 | <1.0E-323 |
| *purL* | SAUSA300_RS05220 | Purines, pyrimidines, nucleosides, and nucleotides | Purine ribonucleotide biosynthesis | -0.797 | <1.0E-323 |
| *brnQ1* | SAUSA300_RS00985 | Transport and binding proteins | Amino acids, peptides and amines | -0.784 | <1.0E-323 |
| *accD* | SAUSA300_RS08990 | Fatty acid and phospholipid metabolism | Biosynthesis | -0.784 | <1.0E-323 |
| *nasE* | SAUSA300_RS12945 | Central intermediary metabolism | Nitrogen metabolism | -0.784 | <1.0E-323 |
| *ebh* | SAUSA300_RS07235 | Virulence | Virulence | -0.782 | <1.0E-323 |
| *nickel ABC transporter substrate-binding protein* | SAUSA300_RS00370 | Transport and binding proteins | Cations and iron carrying compounds | -0.779 | <1.0E-323 |
| *mecA* | SAUSA300_RS00165 | Cell envelope | Biosynthesis and degradation of murein sacculus and peptidoglycan | -0.774 | <1.0E-323 |
| - | SAUSA300_RS07125 | - | - | -0.774 | <1.0E-323 |
| *2-hydroxyacid dehydrogenase* | SAUSA300_RS12450 | Amino acid biosynthesis | Serine family | -0.771 | <1.0E-323 |
| *sodium:proton antiporter* | SAUSA300_RS04570 | Transport and binding proteins | Carbohydrates, organic alcohols, and acids | -0.762 | <1.0E-323 |
| *signal recognition particle sRNA large type* | SAUSA300_RS15100 | - | - | -0.762 | <1.0E-323 |
| *pgm* | SAUSA300_RS04095 | Energy metabolism | Glycolysis/gluconeogenesis | -0.759 | <1.0E-323 |
| *rpoC* | SAUSA300_RS02825 | Transcription | DNA-dependent RNA polymerase | -0.759 | <1.0E-323 |
| - | SAUSA300_RS09010 | Purines, pyrimidines, nucleosides, and nucleotides | Purine ribonucleotide biosynthesis | -0.750 | <1.0E-323 |
| *rpoA* | SAUSA300_RS12020 | Transcription | DNA-dependent RNA polymerase | -0.749 | <1.0E-323 |
| *rpsJ* | SAUSA300_RS12155 | Protein synthesis | Ribosomal proteins: synthesis and modification | -0.746 | <1.0E-323 |
| *clpB* | SAUSA300_RS04730 | Protein fate | Protein folding and stabilization | -0.743 | <1.0E-323 |
| *phage portal protein* | SAUSA300_RS07655 | Mobile and extrachromosomal element functions | Prophage functions | -0.741 | <1.0E-323 |
| *mntB* | SAUSA300_RS03320 | Transport and binding proteins | Cations and iron carrying compounds | -0.737 | <1.0E-323 |
| *dynA* | SAUSA300_RS07265 | Protein synthesis | Other | -0.737 | <1.0E-323 |
| *secDF* | SAUSA300_RS08680 | Protein fate | Protein and peptide secretion and trafficking | -0.736 | <1.0E-323 |
| *parC* | SAUSA300_RS06790 | DNA metabolism | DNA replication, recombination, and repair | -0.736 | <1.0E-323 |
| *polC* | SAUSA300_RS06265 | DNA metabolism | DNA replication, recombination, and repair | -0.736 | <1.0E-323 |
| *ftsH* | SAUSA300_RS02625 | Protein fate; Cellular processes | Degradation of proteins, peptides, and glycopeptides; Cell division | -0.733 | <1.0E-323 |
| *lctP1* | SAUSA300_RS00580 | Transport and binding proteins | Carbohydrates, organic alcohols, and acids | -0.732 | <1.0E-323 |
| *tuf* | SAUSA300_RS02850 | Protein synthesis | Translation factors | -0.728 | <1.0E-323 |
| *pde2* | SAUSA300_RS09005 | Cell envelope | Biosynthesis and degradation of murein sacculus and peptidoglycan | -0.727 | <1.0E-323 |

| | | | | | |
|---|---|---|---|---|---|
| *16S rRNA pseudouridine(516) synthase* | SAUSA300_RS09280 | Protein synthesis | tRNA and rRNA base modification | -0.720 | <1.0E-323 |
| *RNase P RNA component class B* | SAUSA300_RS15340 | - | - | -0.720 | <1.0E-323 |
| *gidA* | SAUSA300_RS14685 | Protein synthesis | tRNA and rRNA base modification | -0.718 | <1.0E-323 |
| *malA* | SAUSA300_RS07950 | Energy metabolism | Sugars | -0.716 | <1.0E-323 |
| *gcvPA* | SAUSA300_RS08170 | Energy metabolism | Amino acids and amines | -0.715 | <1.0E-323 |
| *rpoB* | SAUSA300_RS02820 | Transcription | DNA-dependent RNA polymerase | -0.715 | <1.0E-323 |
| *menD* | SAUSA300_RS05085 | Biosynthesis of cofactors, prosthetic groups, and carriers | Menaquinone and ubiquinone | -0.713 | <1.0E-323 |
| *nupG* | SAUSA300_RS03380 | Transport and binding proteins | Nucleosides, purines and pyrimidines | -0.711 | <1.0E-323 |
| *serine hydrolase family protein* | SAUSA300_RS09740 | - | - | -0.710 | <1.0E-323 |
| *adk* | SAUSA300_RS12045 | Purines, pyrimidines, nucleosides, and nucleotides | Nucleotide and nucleoside interconversions | -0.708 | <1.0E-323 |
| *membrane protein* | SAUSA300_RS07370 | - | - | -0.707 | <1.0E-323 |
| *thiN* | SAUSA300_RS06040 | Biosynthesis of cofactors, prosthetic groups, and carriers | Thiamine | -0.705 | <1.0E-323 |
| *purH* | SAUSA300_RS05240 | Purines, pyrimidines, nucleosides, and nucleotides | Purine ribonucleotide biosynthesis | -0.703 | <1.0E-323 |
| *terminase* | SAUSA300_RS10650 | - | - | -0.703 | <1.0E-323 |

Tables S3-S7 uploaded as Excel files.
